# Amyloid beta oligomers modulate neuronal autophagy through the primary cilium

**DOI:** 10.1101/2021.06.02.446758

**Authors:** Olatz Pampliega, Federico N. Soria, Narayana Pineda-Ramírez, Erwan Bezard

## Abstract

The major neurodegenerative diseases, like Alzheimer’s disease (AD), accumulate neuropathogenic proteins that compromise autophagic function. In AD, autophagy contributes to intracellular APP processing and amyloid beta (Aβ) generation by mutant presenilin-1 (PS1). However, how extracellular soluble Aβ oligomers (Aβo) impact intracellular autophagy is not well understood. The primary cilium (PC), a signaling organelle on the surface of mature neurons and glia, is able to bind Aβ. Since PC signaling pathways knowingly modify autophagy in non-brain cells, we here investigated the role of neuronal PC in the modulation of autophagy during acute extracellular Aβo overload. Our results show that, *in vivo*, recombinant Aβo require the presence of neuronal PC to modulate early autophagy and to induce the accumulation of autophagic vacuoles in an age-dependent manner. We show that activated Akt mediates these effects in an age-dependent manner, and that ciliary p75NTR receptor is required to block autophagy by Aβo. These findings demonstrate that neuronal PC in the adult brain participates in the deleterious effects mediated by soluble Aβo. The PC should therefore be considered as a target organelle to modulate autophagy for the treatment of neurodegenerative diseases.

**Highlights:**

- Aβo requires the neuronal PC to impair learning in young and old mice.
- Autophagy in whole hippocampus differs from autophagy response in hippocampal neurons.
- Aβo induce autophagolysosome accumulation through primary cilia- and age-dependent Akt phosphorylation.

## INTRODUCTION

Aging is the major risk factor for neurodegenerative diseases like Alzheimeŕs disease (AD), where soluble amyloid beta oligomers (Aβo) accumulate in brain regions such as the hippocampus, impairing memory and learning ^1^. Aging also compromises autophagy, the catabolic pathway in charge of degrading deleterious organelles and aggregates. The age-dependent decline in autophagy is widely described ^2^, and the events related with the malfunction of this pathway impact the course of the main neurological pathologies, including AD ^3^.

The generation of new Aβ peptides can happen intracellularly through the processing of the amyloid precursor protein (APP) in autophagic vesicles (AV) ^4^. Mutations in AD related genes such as presenilin-1 (PS-1) have been reported to increase the processing and accumulation of Aβ in AVs, leading to the accumulation of undigested material in the neurites of AD patients ^5–7^. According to the amyloid hypothesis, once generated, Aβ peptides are released to the extracellular space where they may oligomerize ^8^, although a less popular hypothesis posits that oligomerization of Aβ can start inside neurons ^9^.

It has recently been described that, in non-neuronal cells, extracellular Aβ modifies the structure and function of the primary cilium (PC) ^10^, a signaling organelle sitting on the surface of most cell types. Aβ alters the activity of the cilia-dependent Hedgehog (Hh) pathway ^10^, suggesting that cilia-mediated Aβ effects can impact intracellular events. Interestingly, signaling pathways clustered in the PC are able to modify autophagy activity through Hh activation in response to starvation ^11^, whereas other extracellular stimuli such as shear stress modulate autophagy in the kidney by cilia located polycistin 2 ^12^. Thus, diverse extracellular stimuli could impact autophagy activity through different ciliary pathways depending on the cell type and its biological function. However, whether extracellular Aβo are able to modify cilia morphology and activity in adult neurons and potentially affect neuronal autophagy during aging has yet to be discovered.

Here we study whether extracellular soluble Aβo interfere with intracellular autophagy in hippocampal adult neurons through neuronal ciliary pathways, and whether these events are altered during aging. We hypothesize that extracellular soluble Aβo modify autophagic activity in neurons through ciliary signaling pathways, and that these events are differentially regulated in young and old mice.

## RESULTS

### Conditional deletion of the primary cilium in young and aged mice

To investigate whether extracellular Aβo modify intracellular autophagy through ciliary pathways in the adult brain, we generated a Ift88-SLICK-H (Ift88F/F::Thy1-Cre/+) mouse line where we conditionally ablate cilia presence in projection neurons, and thus, we avoid the deleterious effects of cilia depletion during development. Ift88-SLICK-H mice were generated by crossing Ift88 flanked loxP mice (Ift88 F/F) with SLICK-H mice (Thy1-Cre/ERT2,-EYFP). PC was deleted in Thy1^+^ neurons upon intraperitoneal injection of 75mg tamoxifen/kg body weight during 5 consecutive days in mice 3 and 15 months old.

We confirmed the efficiency of Cre-lox recombination in the regions of interest by real-time PCR. Brain extracts from the hippocampus (HIPP) and cortex (CTX) showed a high efficient cre-lox recombination at 3mo and 15mo, as showed by the intensity of the 570pb band. Other regions such as the striatum (STR) or the olfactory bulb (BLB), where Thy1 expression is limited in SLICK-H mice^13^, showed a weaker recombination at 3mo that is increased at 15mo. Absence of Cre-lox recombination was confirmed in brain regions of control mice (F/F), in wild types (WT), and in the tail of F/+ mice where the 570pb is not present (**Supplementary Fig. 1A**).

We then confirmed the depletion of IFT88 protein in hippocampal extracts from 3mo and 15mo male IFT88-/-^Thy1^ and F/F control mice (**Supplementary Fig. 1B**), where levels drop a 58% both at 3mo and 15mo (**Supplementary Fig. 1C**). However we noticed that in male mice IFT88 protein depletion is larger than in females, where levels drop by 76 ± 6.6 % in 3mo and a 60 ± 8.7 % in 15mo IFT88-/-^Thy1^ males (**Supplementary Fig.1B)**, compared to a 42 ± 13% (3mo) and 57 ± 8% (15mo) IFT88 decrease in females in the hippocampus **(Supplementary Figure 1B**).

Decreased IFT88 levels in the hippocampus are accompanied by a depletion of neuronal cilia in hippocampal CA1 region, which is highlighted by immunostaining against adenylyl-cyclase III (ACIII) (**Supplementary Fig. 1D, E**). Number of cilia in CA1 molecular layer is decreased 70.6 ± 9% at 3mo and 66.7 ± 7% at 15mo. Similarly, cilia in aged control mice is 20 % longer than PC in young animals (6.08 ± 1.15 µm at 15mo vs. 5.19 ± 1.19 µm at 3mo), indicating that aging has a trend to increase neuronal cilia length. In contrast, remaining cilia in IFT88-/-^Thy1^ mice show a significant reduction in PC length, being of 2.27 ± 0.48 µm at 3mo, and 3.88 ± 1.18 µm at 15mo (**Supplementary Figure 1E**). Altogether, these data show that peripheral tamoxifen injection efficiently induces cilia deletion in hippocampal Thy1+ neurons.

### Amyloid beta oligomers do not alter neuronal primary cilia structure in vivo

Once we confirmed conditional cilia depletion in the hippocampus of young and old mice, we proceeded to inject Aβo and their control peptide Aβ scrambled (Aβsc) in male and female Ift88F/F (control) and IFT88-/-^Thy1^ mice. Aβo and Aβsc were slowly infused by stereotaxic surgery into the brain ventricles, from where they diffuse into the adjacent parenchyma ^14, 15^. We then analyzed the hippocampus of these mice by fluorescence microscopy and biochemistry, 14 days after Aβo (or Aβsc) inoculation.

As we were interested in deciphering the role of Aβo as a ciliary signaling molecule, we initially wondered whether extracellular Aβo modify ciliogenesis or PC structure. First, immunoblot analysis shows that Aβo do not significantly modify IFT88 levels in the hippocampus (**Figure 1A**), although there is a trend to increased IFT88 protein in Aβo-injected control male mice (29.6 ± 0.3 % increase) that is accentuated in old males (54.3 ± 0.27% increase, **Supplementary Figure 2A**). In contrast, in females, Aβo reduce IFT88 levels at 3mo, whereas this protein levels are unchanged in 15mo females (**Supplementary Figure 2A**).

**Figure 1.**
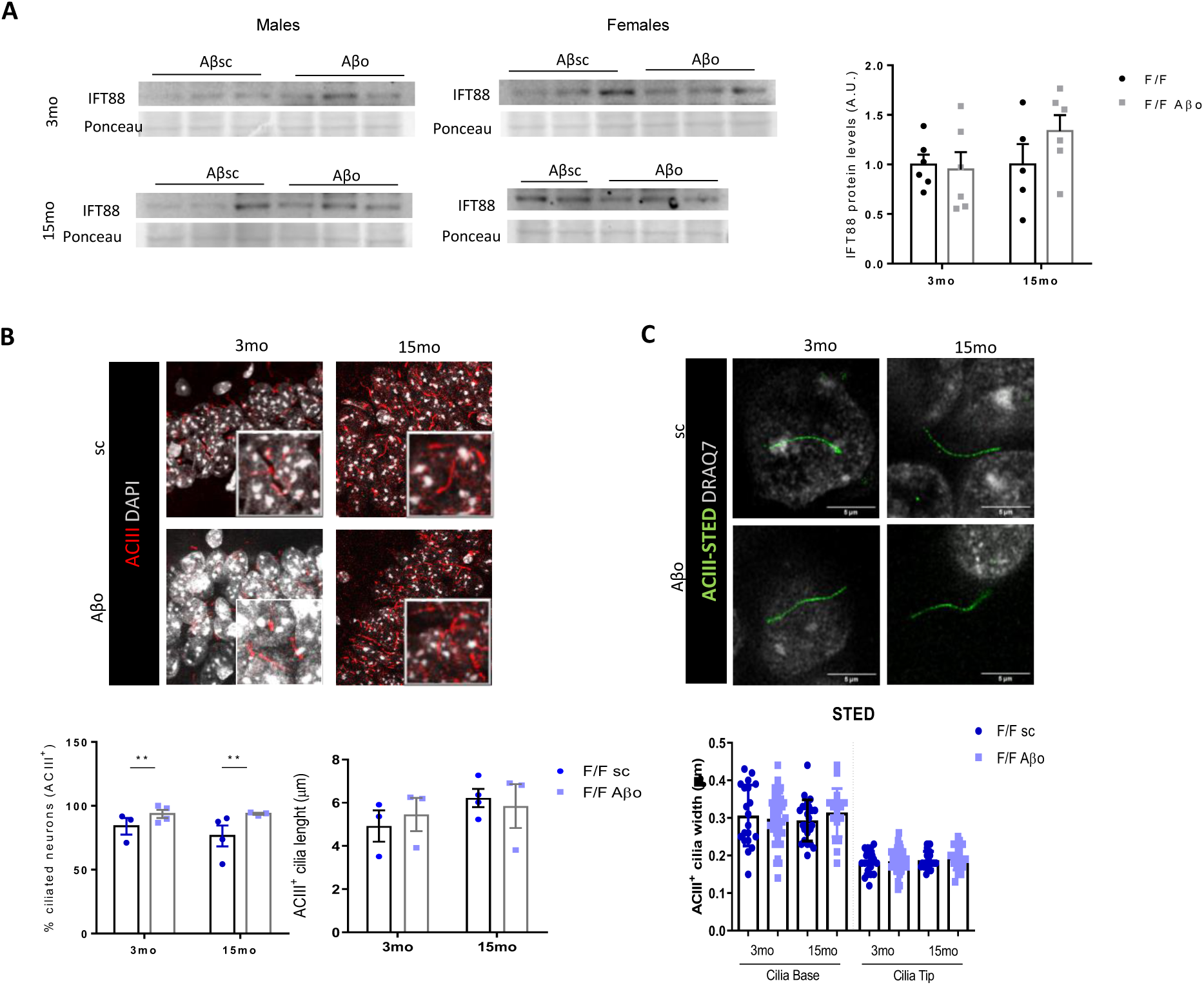
Amyloid beta oligomers do not alter neuronal primary cilia structure *in vivo*. (A) Right; Immunoblot for IFT88 and actin in hippocampus from young (3mo) and old (15mo) F/F male mice injected with either Aβsc or Aβo (left). Left; and densitometry. *n = 3*. (B) Up; Immunohistochemistry for ACIII and DAPI in CA1 from young (3mo) and old (15mo) F/F mice. (E) Down; % of ACIII+ (ciliated) cells in CA1 and cilia length in the same samples. (C) Up; STED microscopy of ACIII+ and DRAQ7 in CA1 from young (3mo) and old (15mo) F/F mice injected with either Aβsc or Aβo. Down; quantification of cilia width at the base and the tip in the same samples; number of quantified cilia are represented. A total of *n = 3-4* hippocampi from 3 different animals per experimental condition are quantified. Mean ± s.e.m is shown. Two-way ANOVA and Sidak’s multiple comparisons test; * *p =* 0.05, ** *p =* 0.01, *** *p =* 0.001.

In addition, confocal imaging of CA1 cilia showed that Aβo increases the percentage of ciliated neurons (a 9.7 ± 6.5% increase at 3mo, and a 17.3 ± 0.7% at 15mo; **Figure 1B**). This increase in ciliated neurons was not accompanied by changes in cilia length between Aβo and Aβsc (**Figure 1B**). Since cilia transversal dimension falls below the diffraction limit of light microscopy, we used STED to analyze this parameter, and we observed no changes in cilia width, as well as no apparent morphological alteration in cilia from Aβo nor Aβsc groups (**Figure 1C**). Altogether our data suggest that acute injection of Aβo does not modify cilia morphology and length in the hippocampus, but increases the number of ciliated neurons.

### Amyloid beta oligomers require the primary cilium to impair learning

Soluble Aβo are thought to be responsible, at least in part, for the memory deficits in AD patients and AD mouse models ^16, 17^. Aβo infusion into the hippocampus leads to cognitive impairment ^14^. In order to ratify the effects of soluble Aβo over memory, we subjected the mice to the Novel Object Recognition test (NOR), which provides and index of recognition memory in rodents by measuring the ability to discriminate between familiar and unfamiliar objects ^18^ (**Figure 2A**). We first noticed that time to reach criterion (i.e. the time needed to actively explore both objects for 20 s in total) was similar in all mice regardless the treatment or genotype, ruling out differences in exploratory activity between these groups. Time to reach criterion in control F/F mice ranged from 202.8 ± 67.75 s in young to 377.1 ± 118.6 s in old, while in IFT88-/-^Thy1^ ranged from 273.3 ± 141.0 s to 362.6 ± 107.3 s (**Figure 2B**). Additionally, there is a trend to higher time to reach criterion in old mice (**Figure 2B**), suggesting that aging might be interfering with their exploratory activity. However this hypothesis needs additional experiments to be confirmed.

**Figure 2.**
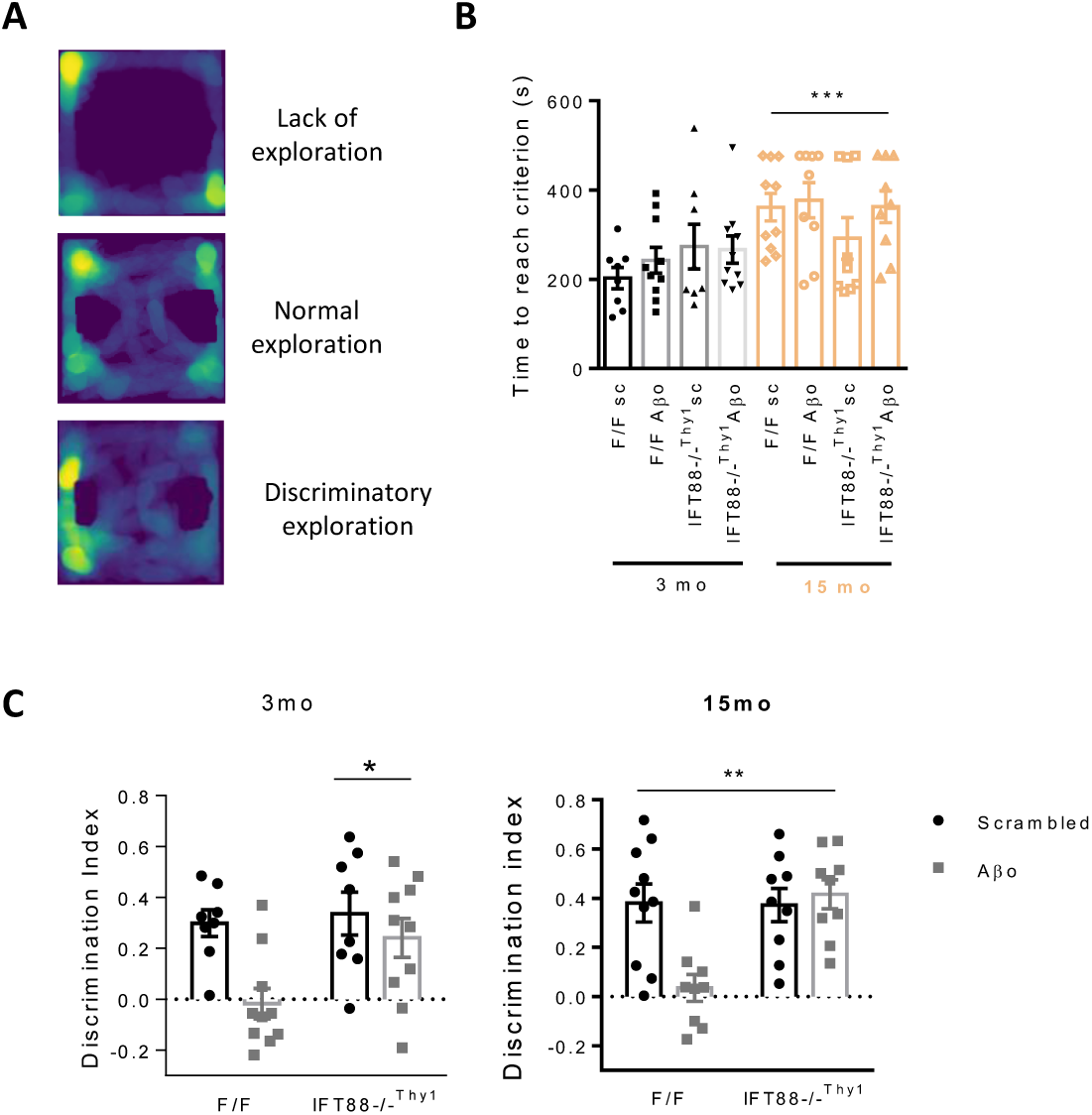
Amyloid beta oligomers require the neuronal primary cilium to impair learning. (A) Cumulative trackings of F/F mice at different phases of NOR experiments. (B) Time to reach criterion in NOR experiment for female and male young (3mo) and old (15mo) F/F and IFT88-/-^Thy1^ injected with either Aβsc or Aβo. (C) Discriminatory Index (DI) in 3mo (left) and 15mo (right) F/F and IFT88-/-^Thy1^ injected with either Aβsc or Aβo. DI= (T_N_-T_F_)/(T_N_+T_F_). *n* = 8-10 female and male mice per experimental condition are analysed. Mean ± s.e.m is shown. Two-way ANOVA and Sidak’s multiple comparisons test; * *p =* 0.05, ** *p =* 0.01, *** *p =* 0.001.

The main readout from the NOR test is the Discrimination index (DI), which indicates the percentage of time spent actively exploring the novel object with respect to the familiar one ^19^, where DI > 1 indicates a discriminatory exploration towards the novel object, D = 0 shows no discrimination and D < 1 represents more time exploring the familiar object. Either 0 or negative values, thus, indicate impaired recognition memory. In consonance with previous reports ^15^, we observed that DI was significantly reduced after Aβo intraventricular injection in control (F/F) mice, indicating that soluble Aβ impairs memory in these animals. Young mice showed a drop in DI from 0.299 ± 0.05 in Aβsc to -0.018 ± 0.05 in Aβo, along with old mice which showed a similar decrease; from 0.380 ± 0.08 in Aβsc to 0.035 ± 0.05 in Aβo (**Figure 2C**). Interestingly, Aβo did not reduce DI in IFT88-/-^Thy1^. Young IFT88-/-^Thy1^ mice showed a DI of 0.336 ± 0.08 upon Aβsc and 0.241 ± 0.08 upon Aβo, while in old IFT88-/-^Thy1^ mice DI was 0.371 ± 0.07 upon Aβsc and 0.416 ± 0.06 upon Aβo inoculation, all of these levels being comparable to F/F control mice (**Figure 2C**). Altogether these data indicate that soluble Aβo need the presence of neuronal PC to impair learning in mice. Of note is that female and male mice did not show sex related differences in learning capacity or in the effects of PC-dependent Aβo cognitive impairment (**Supplementary Figure 2B)**.

### Amyloid beta oligomers activate early autophagy in a cilia- and age-dependent manner

We next studied the autophagy status in hippocampal extracts and fixed tissue from control F/F and IFT88-/-^Thy1^ young and old mice injected with either Aβsc or Aβo. Hippocampal extracts from male and female mice were analysed separately in order to account for sex-related differences in autophagy ^20–22^.

Early steps of autophagy were studied by checking the levels of ATG14, ATG16L1 and ATG7 proteins in hippocampal extracts (**Figure 3A**). ATG14 is a marker for autophagy activation, while ATG16L1 levels reflect the recruitment of early membranes for autophagosome formation. ATG7 is an E1-like activating enzyme required for autophagy. Overall we found that while in male mice Aβo related autophagy was cilia- and age-dependent, female mice did not regulate Aβo-related autophagy through the PC. Thus, we show male related changes in the main figures and we describe changes in females in the supplementary material.

**Figure 3.**
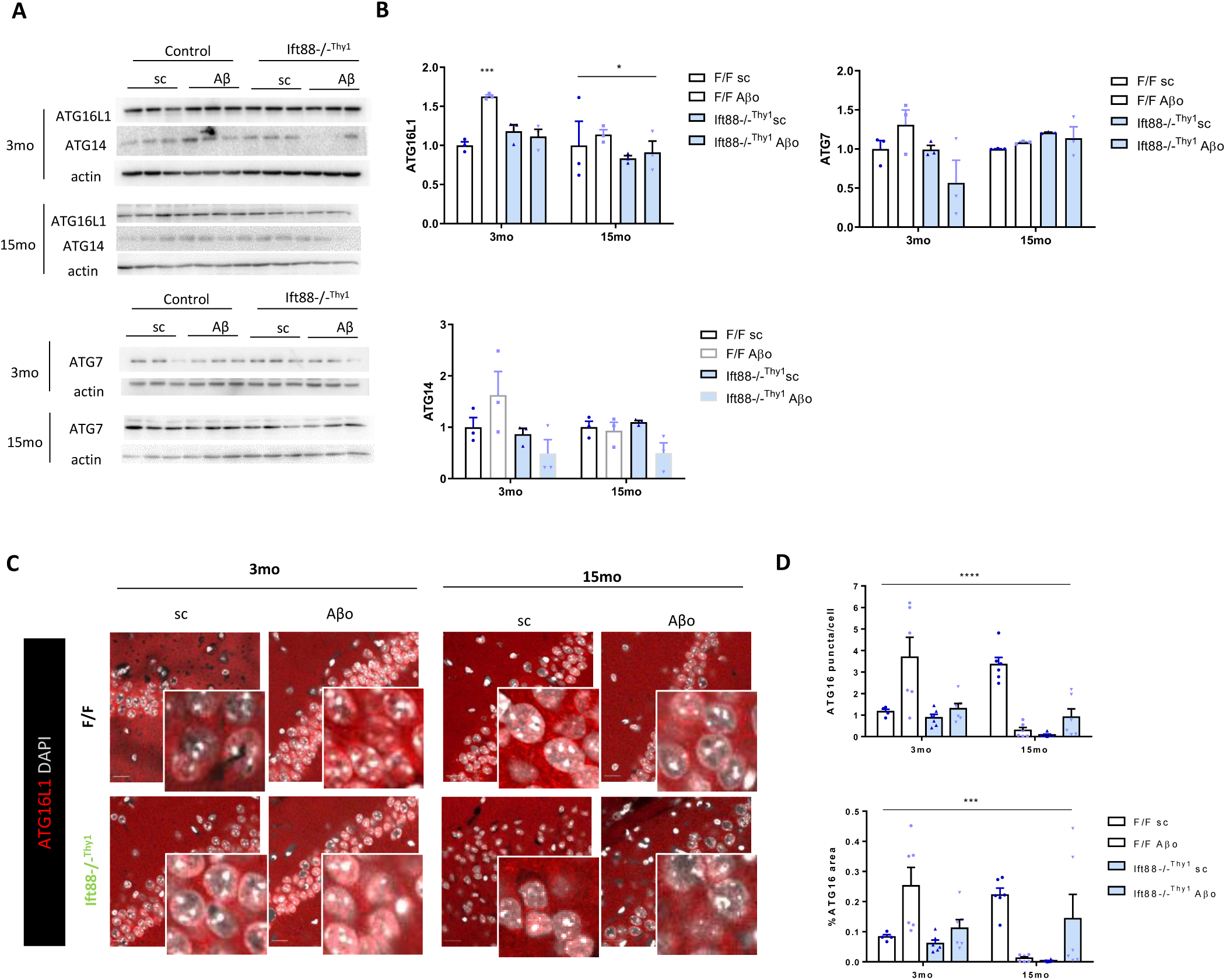
Amyloid beta oligomers activate early neuronal autophagy in a cilia- and age-dependent manner. (A) Immunoblots for early autophagy markers ATG16L1, ATG14, and ATG7, and actin in young (3mo) and old (15mo) F/F and IFT88-/-^Thy1^ male mice injected with either Aβsc or Aβo. (B) Densitometry of immunoblots in (A), *n* = 3. (C) Immunohistochemistry for ATG16L1 and DAPI in CA1 from young (3mo) and old (15mo) F/F and IFT88-/-^Thy1^ mice injected with either Aβsc or Aβo. (D) Quantification of ATG16L1 puncta per cell in CA1 (up) and % of ATG16L1 occupied area in (C). *n* = 6 hippocampi from 3 different mice per experimental condition are analysed. Mean ± s.e.m is shown. Two-way ANOVA and Sidak’s multiple comparisons test; * *p =* 0.05, ** *p =* 0.01, *** *p =* 0.001.

In young F/F males, Aβo induced a slight, non-significant, upregulation of ATG14, and ATG7 proteins. In contrast, ATG16L1 increased upon Aβo in young control F/F animals (62.6 ± 0.21% increase), while in young IFT88-/-^Thy1^ there was no change (+18.08 ± 8.4% in Aβsc and +11.33 ± 9.2% in Aβo). Moreover, these observed changes in ATG16L1 are abolished in old animals (13.93 ± 6.1% increase upon Aβo in F/F), indicating that upregulation of ATG16L1 in response to Aβo might be an age- and cilia-dependent event (**Figure 3C**). Data also indicated that early autophagy is differently modulated in female mice. First, from early autophagy markers, only levels of ATG14 are significantly changed in females, where Aβo reduced the levels of this protein and the lack of neuronal cilia led to a depletion. Aβo also reduced ATG16L1 (20.7 ± 16.4%) in F/F females. In IFT88-/-^Thy1^ young females ATG16L1 was a 26.8 ± 7.9% lower, and a 35.7 ± 4% lower upon Aβo. However, ATG16L1 levels increased in aged females (a 24.3 ± 27% in control and a 56.2 ± 31% in IFT88-/-^Thy1^ females) (**Supplementary Figure 3**). Overall these results suggest that, in contrast to males, females Aβo do not modulate autophagy in a cilia-dependent manner, and thus, implies that autophagy would be differentially modulated in males and females, an important parameters when designing therapeutic strategies targeting cilia-activated autophagy.

Since hippocampal extracts are composed by neurons and glia, it is plausible that neuronal cilia-dependent changes detected by immunoblot are diluted by the presence of protein from other cell types. Therefore, to accurately quantify the expression of early autophagy marker ATG16L1 in neurons from CA1 molecular layer we used fluorescence immunohistochemistry, where ATG16L1 labelling appears as cytoplasmic puncta (**Figure 3C**). In young F/F mice ATG16L1 significantly increase more than three-fold in the Aβo group with respect to Aβsc (1.16 ± 0.11 puncta/cell in Aβsc; 3.69 ± 0.93 puncta/cell in Aβo; 219.43% increase), while IFT88-/-^Thy1^ mice showed a slight non-significant increase from 0.87 ± 0.16 puncta/cell (Aβsc) to 1.29 ± 0.25 puncta/cell (Aβo), indicating that Aβo induces ATG16L1 upregulation in neurons in a cilia-dependent manner in young animals (**Figure 3D**). In contrast, old control F/F mice show increased basal ATG16L1 puncta (3.34 ± 0.33 ATG16L1 puncta/cell in Aβsc, a 189.44% increase compared with young F/F animals) that were abruptly decreased upon Aβo (ATG16L1 puncta/cell decrease to 0.29 ± 0.13). ATG16L1 puncta were also very low in old IFT88-/-^Thy1^ mice (0.07 ± 0.04 and 0.90 ± 0.38 pucnta/cell in old Aβsc and Aβo IFT88-/-^Thy1^ mice, respectively; **Figure 3D**). The fact that ATG16L1 puncta in neurons remains unchanged in IFT88-/-^Thy1^ mice despite the age, suggests that the presence of PC in neurons is a necessary organelle for modulating ATG16L1 levels in these cells.

### Amyloid beta oligomers require the PC to induce autophagolysosome accumulation in neurons

After observing that Aβo upregulate early autophagy in neurons in a cilia-dependent manner, we checked the subsequent steps of the autophagy pathway, i.e. levels of mature autophagosomes and lysosomes, as well as fusioned autophagolysosomes. Immunoblot experiments in hippocampal extracts showed that levels of LC3-II (marker of mature autophagosomes) did not change significantly (**Figure 4A, B**). However, levels of GABARAPL1, an essential protein for later stages of autophagosome maturation ^23^, were significantly increased in old animals upon Aβo injection (a 179.2 ± 84% increase), whereas in young and old IFT88-/-^Thy1^ mice GABARAPL1 levels did not change upon Aβo treatment (**Figure 4C, D**). Regarding lysosomal proteins, LAMP-1 was significantly decreased in IFT88-/-^Thy1^ young mice compared with F/F controls (a decrease of 70.9 ± 1.35% in IFT88-/-^Thy1^ sc, and a 65.6 ± 7.9% in IFT88-/-^Thy1^ Aβo), whereas IFT88-/-^Thy1^ old mice showed increased LAMP-1 hippocampal levels (90.3 ± 4.1% and 55.6 ± 5.6% increase in transgenic sc and Aβo, respectively) (**Figure 4A, B**). Protein levels of LAMP-2 did not change significantly in western blot experiments (**Figure 4A, B**). These data suggest that the PC is required for increasing LAMP-1 labelled organelles in young mice but not LAMP-2+ organelles. Although both LAMP-1 and LAMP-2 are widely used lysosomal markers, both proteins can be heterogeneously distributed in different lysosomal and endocytic organelles ^24^. The cargo receptor p62, which accumulates upon autophagy activation and during aggrephagy, did not show statistically significant changes (**Figure 4C, D**). Data from early autophagy markers and p62 suggest that Aβo activate general autophagy, rather than a specific selective type of autophagy, through ciliary receptors.

**Figure 4.**
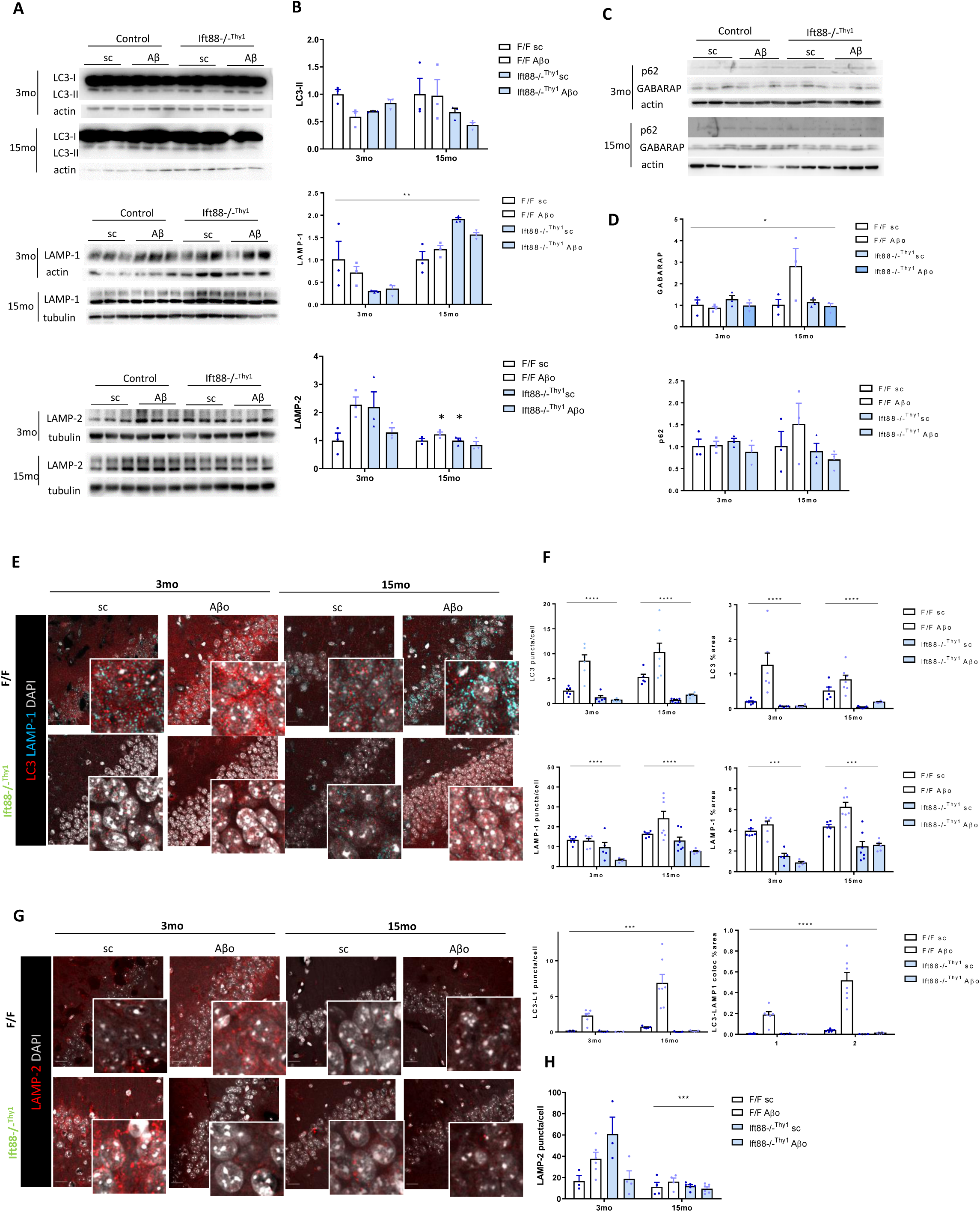
Amyloid beta oligomers require the primary cilium to induce accumulation of autophagolysosomes in neurons. (A) Immunoblot for LC3, LAMP-1, LAMP-2, actin and tubulin in young (3mo) and old (15mo) F/F and IFT88-/-^Thy1^ male mice injected with either Aβsc or Aβo. (B) Densitometry of immunoblots in (A), *n* = 3. (C) Immunoblot for p62, GABARAP, and actin in young (3mo) and old (15mo) F/F and IFT88-/-^Thy1^ male mice injected with either Aβsc or Aβo. (D) Densitometry of immunoblots in (C), *n* = 3. (E) Immunohistochemistry for LC3, LAMP-1 and DAPI in CA1 from young (3mo) and old (15mo) F/F and IFT88-/-^Thy1^ mice injected with either Aβsc or Aβo. (F) Quantification of LC3, LAMP-1 and colocalized LC3-LAMP-1 puncta per cell in CA1 from images in (E). *n* = 6 hippocampi from 3 different animals. (G) Immunohistochemistry for LAMP-2 and DAPI in CA1 from young (3mo) and old (15mo) F/F and IFT88-/-^Thy1^ mice injected with either Aβsc or Aβo. (H) Quantification of LAMP-2 puncta per cell from images in (G). *n* = 6 hippocampi from 3 different animals. Mean ± s.e.m is shown. Two-way ANOVA and Sidak’s multiple comparisons test; * *p =* 0.05, ** *p =* 0.01, *** *p =* 0.001.

To accurately assess the levels of these proteins in hippocampal neurons, we performed double immunohistochemistry for LC3 and LAMP-1 (**Figure 4E**). While LC3 levels reflect the number of autophagosomes, and LAMP-1 the number of lysosomes, particles positive for colocalization of LC3 with LAMP-1 reflect the number of autophagolysosomes, i.e. fused autophagosomes with lysosomes. Immunohistochemistry experiments showed that LC3 levels were significantly increased upon Aβo treatment in a cilia-dependent manner, both in young and old mice. While in young mice LC3 increased from 2.5 ± 0.3 puncta per neuron in Aβsc to 8.5 ± 1.3 in Aβo (241.1% increase), in old mice basal LC3 levels (5.2 ± 0.67 puncta per cell, 108% increase regarding young mice) increased up to 10.25 ± 1.9 puncta per neuron (a 310.9 ± 75.3% increase) upon Aβo (**Figure 4F**). In IFT88-/-^Thy1^ young and old mice LC3 was reduced below basal F/F levels (1.17 ± 0.4 and 0.68 ± 0.14 puncta per cell in young sc and Aβo respectively, and 0.63 ± 0.13 and 1.69 ± 0.19 puncta per cell in old sc and Aβo respectively), indicating that autophagosome modulation occurs through ciliary receptors in neurons. Moreover, old mice show increased basal levels of LC3 (5.21 ± 0.68 puncta per cell), which suggests that either autophagosome biogenesis is increased or these vesicles are not properly mobilized or matured through the autophagy pathway (**Figure 4F**). Immunohistochemistry for the lysosomal marker LAMP-1 showed that in young F/F mice there was no change upon Aβo (13.13 ± 0.8 vs 12.76 ± 1.26 puncta per cell in sc vs Aβo), while in IFT88-/-^Thy1^ mice levels dropped to 9.47 ± 2.7 and 3.23 ± 0.56 puncta per cell upon sc and Aβo, (27.9% and 75.4% decrease), respectively. In old F/F mice, Aβo significantly increased LAMP-1 puncta from 16.22 +/ 0.71 in sc to 24.01 ± 3.8 (59.3% increase) while old IFT88-/-^Thy1^ mice LAMP-1 dropped to similar levels found in young ones (12.84 ± 1.97 and 7.57 ± 0.5 puncta per cell in sc and Aβo respectively). These data suggest that basal and Aβo-induced LAMP-1+ vesicles are regulated through ciliary receptors, an event also observed in old mice (**Figure 4F**). More importantly, we found that Aβo induced a remarkable increase in autophagolysosomes in young F/F (0.12 ± 0.02 to 2.24 ± 0.38 puncta per cell; 1723% increase) and old F/F mice (0.65 ± 0.06 to 6.82 ± 1.26 puncta per cell; 5456.7% increase). Interestingly, autophagolysosome accumulation was abolished in both young and old IFT88-/-^Thy1^ mice (0.07 ± 0.04 in young sc, 0.04 ± 0.01 in young Aβo, 0.03 ± 0.02 in old sc, 0.14 ± 0.03 in old Aβo). Altogether, these data indicate that Aβo induce an accumulation of fused autophagolysosomes in neurons that is enhanced with age, and that these events are dependent on the presence of primary cilia (**Figure 4F**).

We performed immunohistochemistry analysis on LAMP-2, a lysosomal and late endosomal marker, and we found an expression pattern that does not correlate with the results of LAMP-1 (**Figure 4G**). LAMP-2 levels do not show a cilia-dependent pattern, although they are significantly decreased in aged mice (**Figure 4H**).

In contrast to males, female mice did not show that Aβo-mediated autophagy is dependent on the presence of primary cilia in neurons. First, in females LC3-II levels did not show significant changes in hippocampal immunoblots, however LAMP-1 and LAMP-2 hippocampal levels were significantly lower in old females (**Supplementary Figure 4**), although these effects were independent of the genotype and the treatment. Therefore, these data suggest that autophagy is differentially regulated in females, and that lysosomal and endosomal markers are significantly reduced with age, which could result in a poor autophagic activity.

### Phosphorylation of Akt mediates Aβo block of cilia-dependent neuronal autophagy

We show that Aβo activate cilia-dependent autophagy in hippocampal neurons, resulting in the accumulation of non-degraded autophagolysosomes. Moreover, these effects are enhanced with aging, the main risk factor for neurodegenerative diseases. In order to better understand the pathways connecting ciliary signaling with downstream autophagy, we analyzed the hippocampal levels of several signaling pathways at the interface between cilia and autophagy.

First, we studied the expression of phosphorylated AMPK in the hippocampus. AMPK participates in autophagosome maturation and lysosome fusion ^25^, and AMPK activation is coupled to cilia-dependent activation of the Hedgehog pathway ^26^. Moreover, Aβo transiently inhibit AMPK activity in hippocampal neurons ^27^. Thus, we examined phosphorylated AMPK levels by immunoblot (**Figure 5A**) and found that Aβo-induced increase in young F/F mice (68.5 ± 37% increase) is not cilia-dependent (increase of 60.4 ± 27% in IFT88-/-^Thy1^ Aβsc and 25.4 ± 3% in IFT88-/-^Thy1^ Aβo) (**Figure 5B**). Moreover, p-AMPK levels were significantly decreased in old mice (**Figure 5B**), suggesting that aging reduces overall AMPK hippocampal activity.

**Figure 5.**
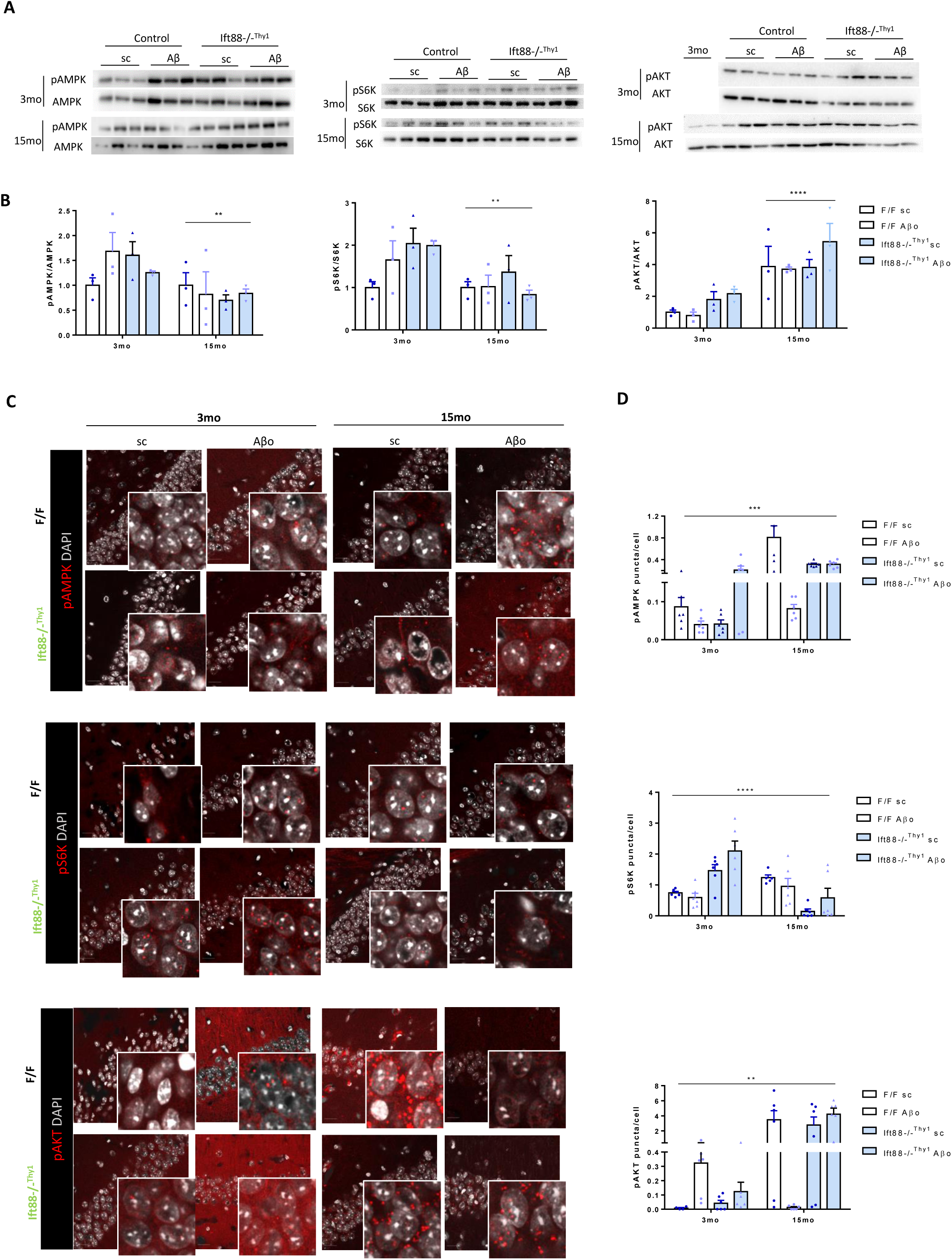
Phosphorylation of Akt mediates Aβo block of cilia-dependent neuronal autophagy. (A) Immunoblots for phospho-AMPK, total AMPK, phospho-P70S6Kinase, total P70S6Kinase, phospho-Akt, total Akt in young (3mo) and old (15mo) F/F and IFT88-/-^Thy1^ male mice injected with either Aβsc or Aβo. (B) Densitometries for immunoblots in (A). (C) Immunohistochemistry for phospho-AMPK, phospho-P70S6Kinase, phospho-Akt, and DAPI in CA1 from young (3mo) and old (15mo) F/F and IFT88-/-^Thy1^ mice injected with either Aβsc or Aβo. (D) Phospho-AMPK, phospho-P70S6Kinase, and phospho-Akt puncta per cell in CA1. *n* = 6 hippocampi from 3 different animals. Mean ± s.e.m is shown. Two-way ANOVA and Sidak’s multiple comparisons test; * *p =* 0.05, ** *p =* 0.01, *** *p =* 0.001.

When we analyzed p-AMPK in hippocampal neurons by immunohistochemistry (**Figure 5C-D**) we found that Aβo induced a decrease in p-AMPK puncta in young F/F mice (from 0.08 ± 0.02 to 0.04 ± 0.01 puncta/cell in Aβo, a 54% decrease) similarly to previously described ^27^. Young IFT88-/-^Thy1^ mice show decreased basal levels of p-AMPK (0.04 ± 0.01 puncta/cell) that increase upon Aβo injection (up to 0.21 ± 0.07 puncta/cell,) indicating that AMPK regulation in young mice is not mediated by neuronal cilia. Old F/F mice showed increased basal levels of p-AMPK (0.81 ± 0.21 puncta/cell, an 834% from young F/F) when compared to young, which decreased upon Aβo(to 0.08 ± 0.1 puncta/cell, a decrease of 840%), again confirming previous research reporting that Aβo transiently inhibit AMPK activity ^27^. However, in old IFT88-/-^Thy^ p-AMPK levels were unchanged after Aβo treatment (0.31 ± 0.02, and 0.31 ± 0.03 puncta/cell respectively).

We next examined mTOR activity, a master regulator of autophagy ^28^. mTOR is also a downstream effector of ciliary receptors, which is regulated by primary cilia ^29^. In the brain, cilia inhibit mTOR activity necessary for a correct ventricle morphogenesis ^30^. To check for mTOR activity, we measured the phosphorylation levels of its downstream serine/threonine kinase P70S6K. Immunoblot analysis of p-P70S6K, showed no significant change in young mice (**Figure 5A**), although there is a trend to increased p-P70S6K levels in IFT88-/-^Thy^ mice (**Figure 5B**). However, old mice show significantly lower levels of p-P70S6K for all experimental conditions and genotypes (**Figure 5B**), suggesting that regulation of mTOR activity in the whole hippocampus is age-dependent. We next studied p-P70S6K levels in hippocampal neurons by immunohistochemistry (**Figure 5C-D**) and we found that young IFT88-/-^Thy^ mice showed increased p-P70S6K puncta in neurons (1.46 ± 0.2 in sc and 2.1 ± 0.32 puncta per cell in Aβo) comparing to F/F, and that this increase was abolished in old IFT88-/-^Thy^ mice (0.14 ± 0.08 in sc, and 0.58 ± 0.3 puncta/cell in Aβo). We also found that similar to p-AMPK, basal p-P70S6K puncta was increased in old control mice (from 0.75 ± 0.05 in young to 1.24 ± 0.09 puncta/cell in old) (66.0% increase). Aβo did not modify p-P70S6K levels, neither in young nor in old mice. Altogether these data indicate that although neuronal primary cilia regulates mTOR activity in an age-dependent manner, it does not modulate Aβo-induced autophagy changes in hippocampal neurons.

Lastly, we examined Akt phosphorylation in hippocampal neurons. Phosphorylated Akt in S473 has been reported to localize at the basal body of primary cilia ^31^. Akt also activates autophagy and it is a central pathway in the clearance of toxic protein aggregates during neurodegeneration ^32^. We therefore hypothesized that Akt could be mediating Aβo-induced autophagy changes through neuronal ciliary pathways. Immunoblot analysis showed that p-Akt levels in the total hippocampus were unchanged upon Aβo treatment in young and old control male mice (**Figure 5A**). However, young IFT88-/-^Thy^ mice showed a trend to increased p-Akt levels (79% and 116% increase in sc and Aβo, respectively). Interestingly, old F/F mice showed significantly increased levels when comparing with young mice for all experimental conditions and genotypes (**Figure 5B**). These data indicate that aging maintains Akt activity in the whole hippocampus. Female mice immunoblot showed that neither Aβo, nor neuronal primary cilia regulate Akt activity (**Supplementary Figure 5**). However, aging significantly increased Akt activation, suggesting that both in males and females aging is a major regulator of Akt in the whole hippocampus (**Supplementary Figure 5**).

We next studied neuronal Akt activity by immunohistochemistry (**Figure 5C-D**). In young hippocampal neurons, Aβo significantly increased Akt phosphorylation (0.008 ± 0.004 and 0.32 ± 0.1 puncta/cell in sc and Aβo respectively), which was abolished in IFT88-/-^Thy^ mice (0.04 ± 0.02 and 0.12 ± 0.07 puncta/cell in sc and Aβo respectively). Interestingly in old F/F mice, p-Akt was increased in basal conditions (3.47 ± 1.21 puncta/cell), and Aβo downregulated p-Akt puncta (up to 0.01 ± 0.006 puncta/cell). Moreover, old IFT88-/-^Thy^ mice maintain high levels of p-Akt that are unresponsive to Aβo treatment (2.77 ± 1.1, and 4.22 ± 0.81 puncta/cell in sc and Aβo respectively). Altogether these data indicate that in hippocampal neurons Aβo increase Akt phosphorylation in a cilia-dependent manner, and that, these effects are inversed in old mice which present high levels of p-Akt that are downregulated upon Aβo also in a cilia-dependent manner.

### Amyloid beta oligomers impair autophagy activity through ciliary p75NTR receptor

Once we verified that Aβo activate Akt through the neuronal primary cilia and that activation of Akt correlates with changes in autophagy, we wanted to decipher the ciliary pathways mediating these effects. Hippocampal neurons express the p75NTR in their PC, and these cilia are modified in 3xAD-transgenic mice, which produce Aβ and tau ^33^. We therefore hypothesized that Aβo might activate ciliary p75NTR to modulate autophagy through Akt.

Treating primary hippocampal neurons with sub lethal doses of Aβo (100nM, **Figure 6A**), we first confirmed that Aβo phosphorylate Akt (105.15% increase) similarly to *in vivo* experiments, and that treatment with LM11A 31, a non-peptide small-molecule ligand of p75NTR that blocks its activity ^34, 35^, prevents Akt phosphorylation (21.9% increase, **Figure 6B**). Aβo also blocked autophagy flux in hippocampal neurons (76% blockage), and this blockage was prevented by blocking p75NTR with LM11A 31 (**Figure 6C**). Basal LC3-II levels were unchanged upon the different treatments (**Figure 6C**). These data indicate that Aβo-mediated changes in autophagy activity and Akt are mediated by the p75NTR.

**Figure 6.**
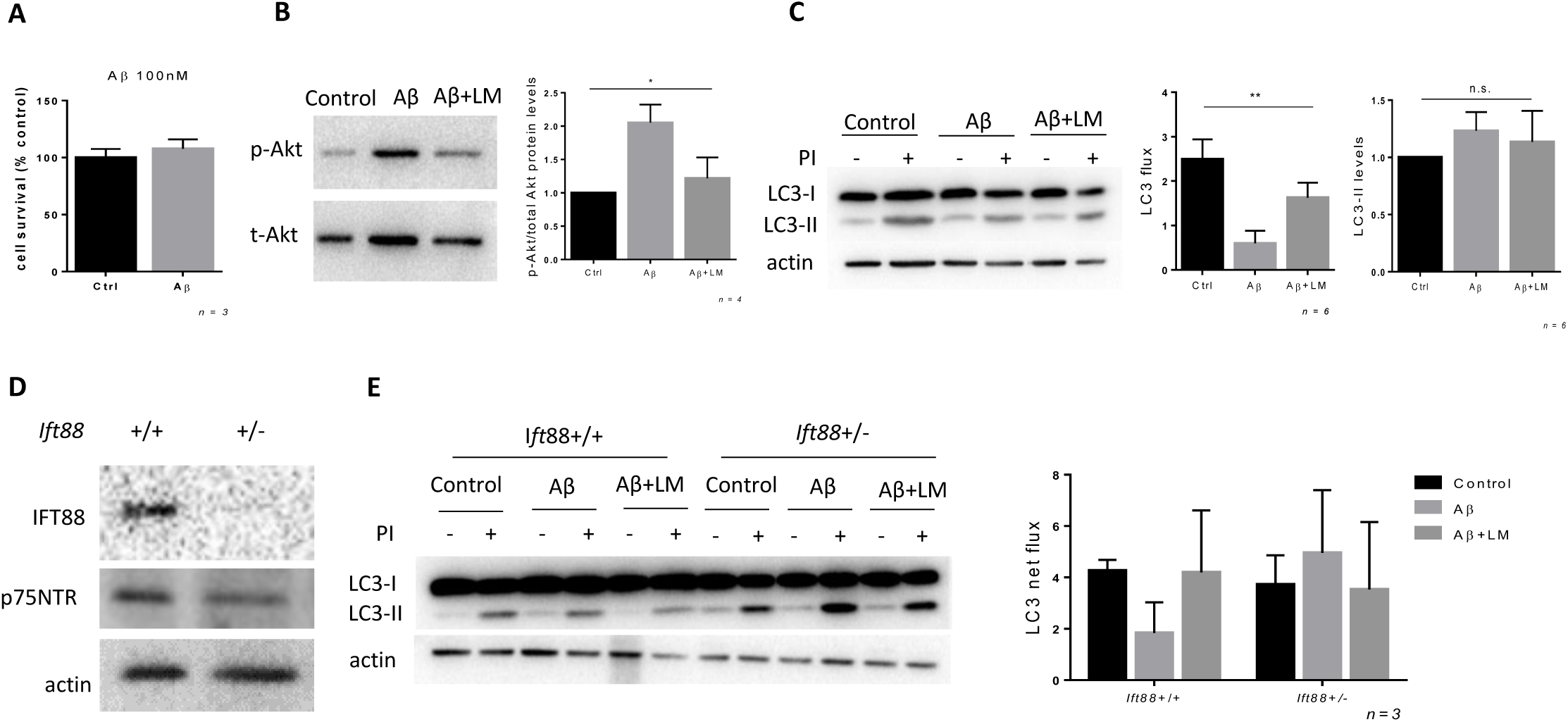
Aβo impair autophagy through ciliary p75NTR. (A) Percentage of survived hippocampal primary neurons treated or not with Aβo 100 nM for 24 h. (B) Left; Immunoblot for phospho-Akt and total Akt in primary hippocampal neurons treated with Aβo 100 nM and LM11A-31 for 24h. Right; Densitometry. *n* = 4. (C) Left; Immunoblot for LC3 and actin in primary hippocampal neurons treated with Aβo 100 nM and LM11A-31 for 24h and incubated in the presence and absence of lysosomal inhibitors (PI) for 1h. Right; densitometry for net autophagy flux and LC3-II levels. *n* = 6. (D) Immunoblot for IFT88, p75NTR and actin in control (+/+) and *Ift88* hemizygous (±) primary hippocampal neurons. (E) Left; Immunoblot for LC3 and actin in primary hippocampal control (+/+) and *Ift88* hemizygous (±) neurons treated with Aβo 100 nM and LM11A-31 for 24h and incubated in the presence and absence of lysosomal inhibitors (PI) for 1h. Right; densitometry for net autophagy flux. *n* = 3. Mean ± s.e.m is shown. Two-way ANOVA and Sidak’s multiple comparisons test; * *p =* 0.05, ** *p =* 0.01, *** *p =* 0.001.

We then tested if p75NTR-dependent changes were mediated by neuronal PC using primary neurons hemizygous for *Ift88*. Lack of *Ift88* in primary neurons is accompanied by a reduction in total levels of p75NTR (**Figure 6D**), suggesting that at least part of this receptor is located in the PC, as previously reported ^36^. We found that loss of *Ift88* in neurons had a trend to prevent autophagy flux blockage by Aβo (4.28 ± 0.4 in +/+ sc; 1.84 ± 1.2 in +/+ Aβo; 4.95 ± 2.4 in ± Aβo; **Figure 6E**). These data suggest that ciliary p75NTR mediates Aβo-induced autophagy blockage in hippocampal neurons.

## DISCUSSION

Recent studies report that extracellular Aβ modifies primary cilia structure and function ^10^ and that hippocampal cilia is shorter in AD animal models ^33^. In addition, it has been widely reported that autophagy malfunctions in AD, with Aβ being generated in acidic vesicles and with AD patients showing accumulation of undigested fused autophagolysosomes ^5^. However how extracellular Aβ could modify intracellular autophagy activity is not well understood. Here we report a novel mechanism whereby extracellular Aβo act through p75NTR ciliary receptors to mediate Akt phosphorylation and thus, to block autophagy activity in hippocampal neurons. We describe that aging reverts Aβo-mediated cilia-dependent Akt signalling, and consequently modifies autophagy outcome in hippocampal neurons.

To understand the signaling role of Aβ we have set up experimental conditions where soluble, non-accumulating Aβo are injected in the brain ventricles of mice, avoiding inflammation ^14^, and cell and cilia damage in the hippocampus. Maintaining an intact cilia structure, absence of cell damage and inflammation allows to study the acute effects of Aβo as signaling molecules, and thus, permits to understand the biochemical basis of Aβo-mediated pathology in the brain, while maintaining the Aβo blocking of recognition memory ^14^.

We report that Aβo impairment of recognition memory is mediated by neuronal primary cilia, regardless animal age or sex. Our data suggest that acute, short-term Aβo effects are mediated by signaling receptors located on the cilium, and also by downstream signaling, rather than by remodeling of synaptic arborization or axonal morphology ^37^. This hypothesis is sustained by the fact that cilia-deficient mice are protected from Aβo-mediated memory impairment, and therefore suggest that cilia-deficient mice maintain the proper synaptic arborization necessary for a correct learning. It is worth noting that in studies where cilia have been removed for longer periods, cilia-defective neurons present shorter dendritic arborizations, corresponding to an immature spine development ^37^, and thus, directly linking the presence of neuronal cilia to a correct dendritic spine formation and functioning in cognition processes.

Although still at very preliminary stage, growing evidence sustain a role for neuronal cilia in AD. *In silico* analysis of cilia interactome shows a significant overlap with AD and an enrichment in neuronal pathways such as neuron maturation or synaptic growth ^38^. Serotonin receptor 5-HT6, a potential target for cognitive improvement in AD ^39^, is mainly located at neuronal PC, and it regulates axonal length also in APP/PS1 mice^40^. The p75NTR, which binds Aβ ^41^, is located in the cilium of hippocampal neurons ^36^ and is reduced in 3xTg-AD mice ^33^. Thus, neuronal PC is emerging as a key organelle mediating AD-related pathology and memory impairment.

In addition to its role in memory performance, we report that extracellular Aβo require the primary cilium to modify autophagy function in hippocampal neurons. We show that the modulation of autophagy in hippocampal neurons is Aβo-, age- and cilia-dependent in male mice, while in females, aging appears to be the sole factor that modifies total hippocampal autophagy significantly. Therefore, different factors modify autophagy in males and females, which should be considered when designing therapeutic strategies. Regulation of autophagy has been reported to be differentially regulated in each sex ^21^, as for example in neurons exposed to nutrient starvation ^22^. Congdon also proposes that male and female differences in autophagy are due to overall lower autophagy in females and to the effect of sex hormones, which contribute to higher frequency of AD in women ^20^. In this line, we show that total hippocampal autophagy decreases in old females, in contrast to males where hippocampal neurons regulate autophagy through ciliary signaling. Sex related differences in AD also include the fact that women with higher Aβ are more prone to cognitive decline than men ^42^ although Aβ levels are similar in men and women with AD ^43^.Thus, increasing evidence point out that targeting autophagy as a therapeutic strategy in AD should consider sex related differences in autophagy regulation.

We report also that aging strongly modulates cilia-dependent autophagy, especially the initation steps that are critical for phagophore recruitment. While in young animals ATG16L1 increase upon Aβo and correlates with higher autophagy fusion and blockage of autophagic flux, in old animals we observe upregulated basal ATG16L1, which is suppressed upon Aβo, indicating that aging modulates the signaling pathways that control ATG16L1. Moreover, the fact that ATG16L1 puncta in neurons remains depleted in cilia deficient mice despite the age, suggests that the pathways controlling ATG16L1 are located within the PC. It has been previously shown that activation of ciliary hedgehog signaling induces ATG16L1 recruitment to ciliary structures ^11^.

In our experimental conditions aging also disrupts the connection between early autophagy and the subsequent steps of autophagosome formation and fusion with lysosomes. In young animals Aβo activate autophagosome formation and induce the accumulation of autophagolysosomes in a cilia-dependent manner. However, in old mice Aβo are unable to activate early autophagy but there is an increase of LC3 puncta (although they occupied less area, suggesting less autophagic capacity), together with an accumulation of autophagolysosomes, which is also cilia dependent. This could be the result of the endocytosis of Aβo and accumulation in late endosomes and lysosomes, which show enlarged morphology ^44^. As depletion of PC disturbs the endocytic capacity at the periciliary area, cilia loss in neurons could prevent Aβo uptake ^45^. In addition, dystrophic axons of AD models accumulate post-TGN LAMP-1^+^ vesicles, which have poor degradative capacity but carry lysosomal components towards the neuronal soma ^46^. We propose that inability to activate early autophagy and autophagosome formation in old animals leds to the accumulation of autophagolysosomes, that could be linked to problems in lysosomal trafficking or enhanced endocytosis of extracellular Aβo ^47, 48^. These options have been previously been reported in AD models, where aging is a major contributor ^5^.

We also show that upstream of autophagy, Aβo promote Akt phosphorylation in a cilia-dependent manner in young animals. Phosphorylated Akt is present at the basal bodies ^31^, thus it is a suitable mediator between ciliary and cytosolic signaling pathways. We show that *in vitro* Akt phosphorylation by Aβo is p75NTR-dependent and blocks autophagy activity. It is worth noting that in our experimental approach Aβo concentration does not interfere with cell viability, and thus, sub-lethal Aβo activate autophagy through Akt phosphorylation ^32^. In contrast, we show that old animals present high basal Akt phosphorylation in hippocampal neurons that is reduced upon Aβo injection. Previous works show that Akt phosphorylation is increased in the aged brain ^49, 50^, and in aged AD models ^50^, although there are regional differences in Akt phosphorylation in the hippocampus ^51^.

It is also worth noting that in young animals, loss of neuronal cilia results in mTOR activation, while in old IFT88-/-^Thy^ mice mTOR activity is decreased. In both cases changes in mTOR are independent of Aβo, and thus, we conclude that in our experimental conditions, Aβo-induced hippocampal neuronal autophagy is mTOR-independent.

In summary our results demonstrate that Aβo require the primary cilium from adult hippocampal neurons to modulate autophagy. First we show that Aβo require the presence of the primary cilium in neurons to impair recognition memory, regardless sex and age. Moreover, we show that Aβo block autophagy activity through ciliary p75NTR and Akt signaling in young mice. We also provide information that old mice present high levels of Akt activity which are suppressed upon Aβo in a cilia-dependent manner, and that also impair autophagy function. Thus, the neuronal primary cilium emerges as a key organelle in transducing Aβo signaling, and therefore, future therapeutic strategies should consider this organelle for new AD therapies.

## METHODS

### Animals

Ift88-SLICK-H (Iftt88F/F::Thy1-Cre/+) were generated by crossing Ift88 flanked loxP mice (B6.129P2-Ift88^tm1Bky^/J from The Jackson laboratory, Stock No: 022409|Ift88^fl^) with SLICK-H mice (STOCK Tg(Thy1-cre/ERT2,-EYFP)HGfng/PyngJ from the Jackson laboratory, Stock No: 012708|SLICK-H). Animals were maintained under standard conditions with food and water *ad libitum* in a normal light/dark cycle. All experimental procedures were approved by the Committee on Animal Health and Care of the University of Bordeaux and French Ministry of Agriculture and Forestry (authorization numbers, APAFIS#8430-2017010412591287 and APAFIS#8719-2017012411024773). Maximal efforts were made to reduce the suffering and the number of animals used. Both male and female mice were used in the experiments.

### Tamoxifen injection and Cre-Lox recombination assay

Tamoxifen (Sigma, #T5648) was dissolved in corn oil at a concentration of 20 mg/ml by shaking overnight at 37 °C. To induce cre-lox recombination, mice were injected intraperitoneally with 75 mg tamoxifen per kg of body weight during 5 consecutive days. Recombination of the LoxP sequences and conditional gene deletion in the target regions, especially hippocampus, was confirmed by PCR as previously described ^52^. Briefly, brain regions (cortex, hippocampus, striatum, olfactory bulb) were collected from 3mo at 15mo mice after sacrifice, and DNA was isolated (Purelink genomic DNA kit, Invitrogen). A piece of tail was used as negative control. qPCR of the target regions was performed for Wt allele, 380pb; Flox allele, 430pb; and Cre-loxP recombined allele, 580pb. Efficient loxP recombination in hippocampus of 3mo and 15mo mice is shown in **Supplementary Figure 1A**.

### Aβo injection

Seven days after the last tamoxifen injection, 3mo and 15mo male and female animals were intraventricularly injected with Aβo and Aβsc. The amyloid beta peptide (Aβ; Amyloid β-Protein (1-42), Bachem #H-7442.0500) and the scrambled control (Aβsc Amyloid β-Protein (1-42)(scrambled) trifluoroacetate salt, Bachem #H-7406.0500) were prepared following manufacturer instructions. For oligomers (Aβo) and Aβsc preparation, HFIP-treated alicuots were resuspended in 2mM anhydrous dimethyl sulfoxide and bath sonicated for 15 min. Samples were then dissolved in 95 µl ice-cold PBS, immediately vortexed for 30 s, and incubated at 4°C for 24 h. This protocol has been previously validated *in vivo* ^14^.Aβo and Aβsc were injected intracerebroventricularly (ICV) to model AD ^53^, as previously described ^14^. A 30-gauge Hamilton syringe coupled to a UMP3T-1 pump (WPI, USA) was used to inject 4 µl of Aβo or Aβsc (infusion rate = 0.5 µl/min) using the following stereotaxic coordinates: anteroposterior (AP) relative to bregma, − 1.0 mm; lateral (L) to midline, 1.3 mm; ventral (V) from the skull surface, − 2.0 mm. Mice were kept under deep isoflurane anaesthesia and buprenorphine (0.1 mg/kg) analgesia during surgery. Animals were weighted and monitored daily during the first week post-surgery.

### Novel Object Recognition

Two weeks after ICV injection of Aβo or Aβsc, recognition memory was assessed in male and female mice. The protocol for Novel Object Recognition (NOR) test has been thoroughly described elsewhere ^19^. Briefly, mice habituation to the test box, was performed for 2 consecutive days, and 24h later, mice were exposed during 8 min to 2 identical objects (Familiarization). The test phase was done 24h later, by replacing one of the familiar objects with a new one, different in shape and texture, and recording in video their exploration and interaction with both objects during 8 min. Time spent actively exploring the objects (i.e. nose less than 1 cm apart and head directed towards the object) was manually measured, until a criterion of 20 s of total object exploration was achieved or the 8 min total time expired. Discrimination index (DI) was calculated as the difference between the time spent actively exploring the novel (N) and the familiar (F) object, divided by the total time exploring the two objects. A higher DI is indicative of a better memory performance of the animal.

### Immunoblot

After euthanasia, hippocampal brain regions for protein analysis were dissected and snap frozen in dry ice. Protein was extracted by mechanical homogenization and subsequent membrane lysis using ice-cold RIPA buffer (Sigma) with freshly added proteinase and phosphatase inhibitors (Roche). After centrifugation of samples at 16,000 xg for 15 min at 4° C, protein concentration was quantified by BCA method (Pierce, Thermo). 10 µg of protein per sample were boiled for 5 min and size separated by electrophoresis in polyacrylamide gels (Bio-Rad). Proteins were transferred over-night at 4 °C to nitrocellulose membranes (Bio-Rad) in 15% methanol transfer buffer. Protein transference to the membrane was checked in 0.1% Ponceau S in 1% acetic acid. Membranes were blocked using 5% non-fat dry milk in TTBS at room temperature (RT°). Primary antibodies were incubated over-night, rocking at 4°C at 1:1000 in a solution of 3% bovine serum albumin (BSA) in TTBS. Primary antibodies for immunoblot included: IFT88 (#13967-1-AP, Proteintech), actin (#A2228, Sigma), ATG16L1 (#PM040, MBL), ATG14 (#PD026, MBL), ATG7 (#AHP1651T, Serotec), LAMP-1 (#AB_528127, Hybridoma Bank), LAMP-2 (#AB_2134767, Hybridoma Bank), tubulin (#T5168, Sigma), LC3B (#NB100-2220, Novus), Phospho-AMPKa Thr172 (#2535, Cell Signaling), AMPK (#2603, Cell Signaling), Phospho-p70 S6 Kinase (Thr389) (#9205, Cell Signaling), p70 S6 Kinase (#9202, Cell Signaling), Phospho-Akt Ser473 (#9271, Cell Signaling), Akt (#9272, Cell Signaling), p75NTR (#AB1554, Millipore). Secondary antibodies conjugated to HRP (Goat anti-Mouse IgG, goat anti-Rabbit IgG, goat anti-Rat IgG from Jackson) were incubated at 1:1000 for 1h at RT° and washed 3 times in TTBS. After, ECL (Pierce) was added and membrane chemiluminiscence was detected (ChemiDoc, Bio-Rad). Chemiluniscence signal from non-saturated images was analyzed using FIJI.

### Immunostaining

Brain tissue for immunohistochemistry was obtained after intracardial perfusion of mice with 4% paraformaldehyde (PFA) in 0.1 M phosphate buffer (PB). Brain were collected, post-fixed in 4% PFA for 24h and dehydrated in 20% sucrose for another 24h. Tissue was subsequently frozen for 2 min in -70°C isopentane and stored at -80°C until further use. Brains were sectioned in cryostat and slices stored in cryopreservation solution (30% glycerol, 30% ethylenglycol in PB) at -20°C. For immunohistochemistry, free-floating sections were first washed in PBS and blocked for 1h at RT° with 5% normal goat serum (NGS), 0.1% Triton-X in PBS. Primary antibodies were diluted 1:200 in 5% NGS, 0.1% tween in PBS and incubated over-night at 4°C. Primary antibodies for immunostaining included: Adenylate cyclase III (#sc-588, Santa Cruz), ATG16L1 (#PM040, MBL), LC3B (#NB100-2220, Novus), LAMP-1 (#AB_528127, Hybridoma Bank), LAMP-2 (#AB_2134767, Hybridoma Bank), Phospho-AMPKa Thr172 (#2535, Cell Signaling), Phospho-p70 S6 Kinase (Thr389) (#9205, Cell Signaling), Phospho-Akt Ser473 (#9271, Cell Signaling). After 3 PBS washes, fluorescent secondary antibodies (1:200 in 5% NGS, 0.1% tween in PBS) were incubated for 2h at RT° and washed 3 times. Secondary antibodies included: Goat anti-rabbit IgG Alexa Fluor 594 (#A-11037, ThermoFisher), Goat anti-rat IgG Alexa Fluor 594 (#A-11007, ThermoFisher), Goat anti-rat IgG Alexa Fluor 680 (#A-21096, ThermoFisher). Hoechst 33258 (ThermoFisher) nuclear staining was added to the first PBS wash after the last secondary antibody. Sections were mounted, dried and coverslipped with Mowiol/DABCO embedding media and left overnight in darkness. For double immunohistochemistry, primary and secondary antibody labelling was done sequentially.

### Confocal and STED microscopy

Confocal images were obtained in a Leica TCS SP8 microscope, maintaining image acquisition settings (laser power, AOTF, detection parameters) between sessions. Image stacks (pixel size ∼100 nm, z-step 0.35 µm) were acquired with 63X Plan Apo CS objectives with oil immersion.

For super-resolution STED microscopy of primary cilia, anti-rabbit conjugated with Alexa-488 Plus (Invitrogen) was used as secondary antibody and nuclei stained with DRAQ5 or DRAQ7. A 595 nm laser was used for depletion of Alexa-488 Plus. STED images were acquired with a 100X objective (numerical aperture = 1.4) with oil immersion at a pixel size ∼20 nm.

### Image analysis

Quantification of acquired images was done in FIJI ^54^. Percentage of ciliated neurons was calculated quantifying the number of cilia presence over the total number of CA1 molecular layer nuclei. Cilia length was quantified in the same region using the measure tool on FIJI. Cilia width was determined at the nanoscale as the full width half maxima (FWHM) of the fluorescence profile traced transversally at the initial and final segments of each cilium.

For puncta quantification as well as for LC3+LAMP-1 puncta colocalization in CA1 molecular layer, we used a custom script (modified from *lyso_puncta_coloc.ijm*, available at https://github.com/SoriaFN/Lysosome_analysis). Briefly, single z-planes were denoised using a difference of Gaussians filter and segmented manually to delineate cells. Puncta was quantified within ROIs by automatic thresholding followed by binaryzation (puncta smaller than 10 px were discarded) and *Analyze Particles* function. Colocalization was obtained from LC3 and LAMP-1 binary images with the ImageCalculator plugin (function “AND”).

Images were processed with Huygens software (SVI) for presentation purposes.

### Primary neuronal cultures

P0-P1 postnatal pups from wild-type and Ift88^F/+^::SLICK-H^Cre/+^ mice were used for primary neuronal cultures. Briefly, after decapitation hippocampi were extracted and enzymatically and mechanically digested in Hank’s buffer. Isolated cells were cultured in Neurobasal media containing B27, Glutamax and 1% psf, and maintained at 37°C in 5% CO_2_. To induce Ift88 deletion in Ift88^F/+^::SLICK-H^Cre/+^ neurons, prior to the cell culture we selected EYFP+ brains under inverted microscope, and at DIV10 we treated hippocampal neurons with 1 µM 4-OH-tamoxifen (Sigma) to induce partial Ift88 deletion. Efficient IFT88 depletion was confirmed by immunoblot (**Figure 6D**). Experiments were performed at DIV14-DIV18.

### Cell survival

LDH release in the culture media from primary hippocampal neurons was measured following manufacturer instructions (Cytotoxicity Detection Kit^Plus^ –LDH, #04 744 926 001, Roche).

### Autophagy flux assays

To block lysosomal proteolysis, primary neuronal cultures were treated or not with ammonium chloride and leupeptine for 2h, and autophagic flux was measured by immunoblot as changes in levels of LC3-II upon inhibition of lysosomal proteolysis (net flux)^55^.

### Statistical analysis

Statistical analysis was carried out in GraphPad Prism v.6.01. For behavioral experiments, immunoblots and immunohistochemistry experiments, two-way ANOVA was applied. All values are given as means +- s.e.m.

### Data availability

The data that support the findings of this study are available from the corresponding authors upon reasonable request. Source data are provided with this paper.

### Code availability

The custom script used for puncta quantifications is available at https://github.com/SoriaFN/Lysosome_analysis.

## ACKNOWLEDGEMENTS

We thank Guillaume Dabée, Melissa Deshors, and Elisabeth Normand for animal care, and Laura Escobar for technical assistance in confocal and STED microscopy. This work was supported by grants from Chairs Junior IdEx Bordeaux, Fondation LECMA Vaincre Alzheimer (FR-15075p), IBRO Return Home Program 2017, Spanish Ministry of Science and Innovation (RTI2018-097948-A-I00) to O.P, Region Nouvelle Aquitaine, Agence Nationale de la Recherche (ANR-16-COEN-0002, ANR-14-OHRI-000L-01), and The Simone and Cino Del Duca Prize from the French Academy of Sciences to E.B. O.P. is a Ramón y Cajal research fellow (RYC-2016-20480), F.N.S. is a Juan de la Cierva-Incorporación postdoctoral fellow (IJCI-2017-32114), and N.P-R. is a graduate student. University of Bordeaux, CNRS, Achucarro Basque Center for Neuroscience and the University of the Basque Country provided infrastructural support. Confocal and STED microscopy were performed at the Imaging Facility of Achucarro Basque Center for Neuroscience.

## AUTHOR INFORMATION

O.P. conceived, designed, performed most experiments, and interpreted the results. F.N.S. performed mouse surgeries, analysed NOR data, assisted with STED imaging and developed image analysis tools. N.P-R. performed biochemical experiments. O.P. wrote the manuscript with input from all authors, secured funding and coordinated the project. E.B. provided funding and revised the manuscript.

## SUPPLEMENTARY FIGURE LEGENDS

**Supplementary Figure 1.**
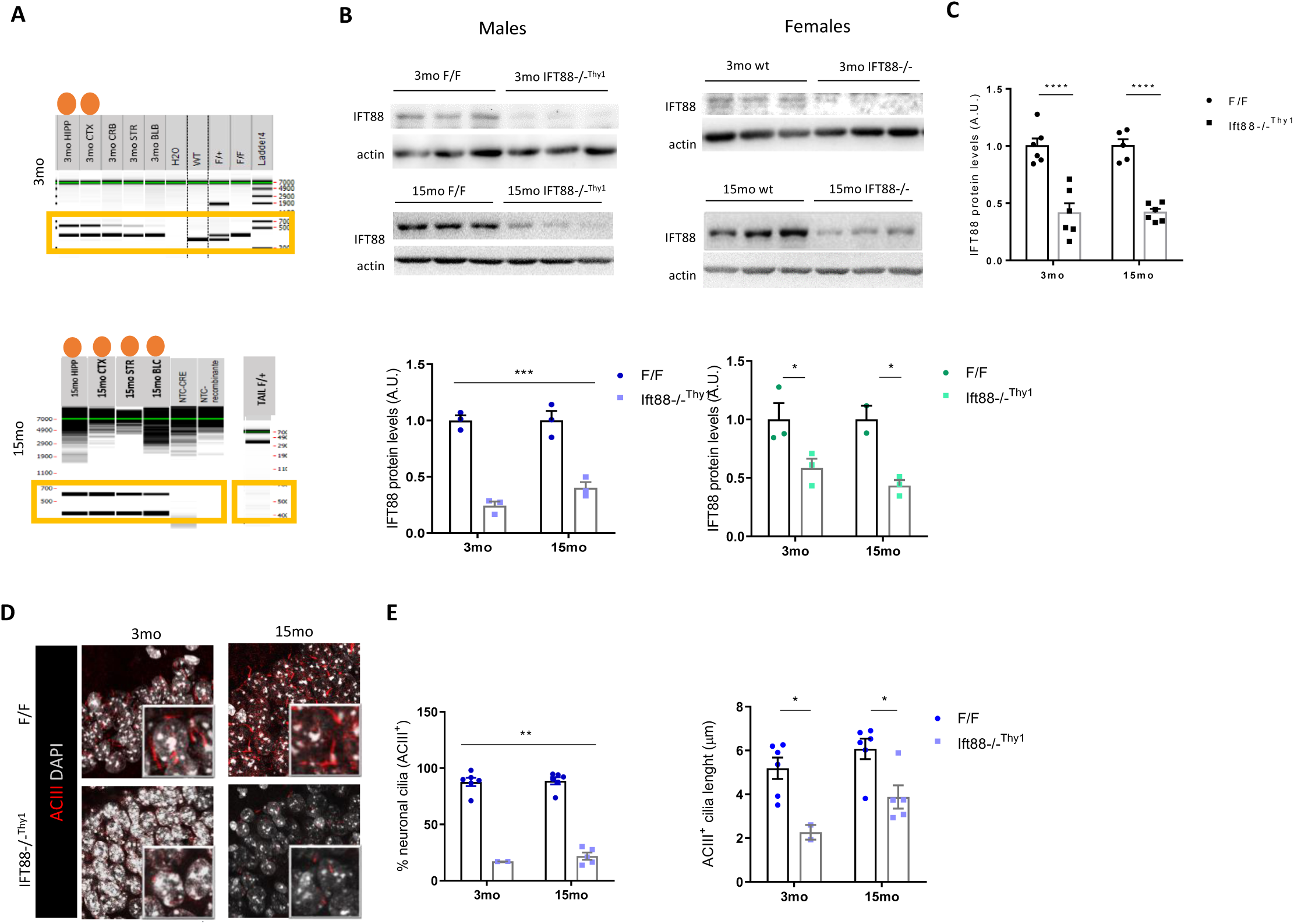
Tamoxifen injection induces Ift88 deletion and cilia loss in the hippocampus. (A) Hippocampal (HIPP), cortex (CTX), cerebellum (CRB), striatum (STR), olfactory bulb (BLB) brain regions subjected to RT-PCR and capillary electrophoresis after DNA isolation to confirm Cre-lox recombination in the target regions. Successful cre-lox recombination shows a band of 570pb. (B) Left; Immunoblot for IFT88 and actin in hippocampus from control and IFT88-/-^Thy1^ young (3mo) and old (15mo) male mice injected with either Aβsc or Aβo (up) and densitometry quantification (down) *n = 3*. Right; Immunoblot for IFT88 and actin in hippocampus from control and IFT88-/-^Thy1^ young (3mo) and old (15mo) female mice injected with either Aβsc or Aβo (up) and densitometry (down) *n = 3*. (C) Quantification of total IFT88 protein levels in male and female mice. (D) Immunohistochemistry for ACIII and DAPI in CA1 from young (3mo) and old (15mo) F/F and IFT88-/-^Thy1^ mice. (E) Quantification of images from (D); Left, % of ACIII+ (ciliated) cells in CA. Right, Cilia length in the same samples. A total of *n = 6* hippocampi from 3 different animals per experimental condition are quantified. Mean ± s.e.m is shown. Two-way ANOVA and Sidak’s multiple comparisons test; * *p =* 0.05, ** *p =* 0.01, *** *p =* 0.001.

**Supplementary Figure 2.**
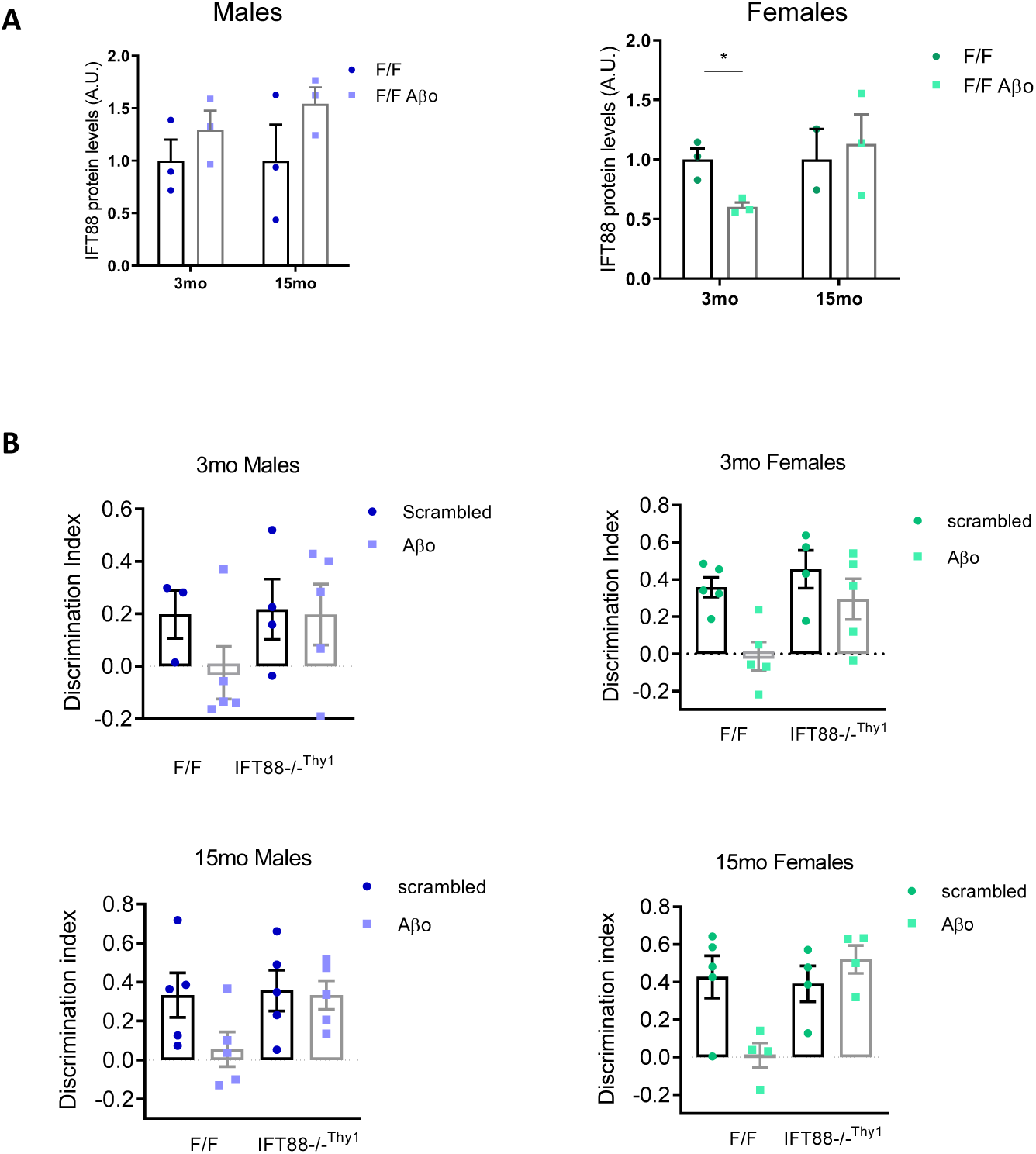
Sex differences in NOR. (A) Right; Quantification of the male immunoblot for IFT88 and actin in young (3mo) and old (15mo) F/F mice injected with either Aβsc or Aβo, from Figure 1A (left) *n* = 3. Left; Quantification of the female immunoblot for IFT88 and actin in young (3mo) and old (15mo) F/F mice injected with either Aβsc or Aβo, from Figure 1A (middle) *n* = 3. (B) DI from NOR experiments expressed in young (3mo) and old (15mo) males and females separately. *n* = 3-5 female or male mice per experimental condition are analysed. Mean ± s.e.m is shown. Two-way ANOVA and Sidak’s multiple comparisons test; * *p =* 0.05, \*\**p =* 0.01, *** *p =* 0.001.

**Supplementary Figure 3.**
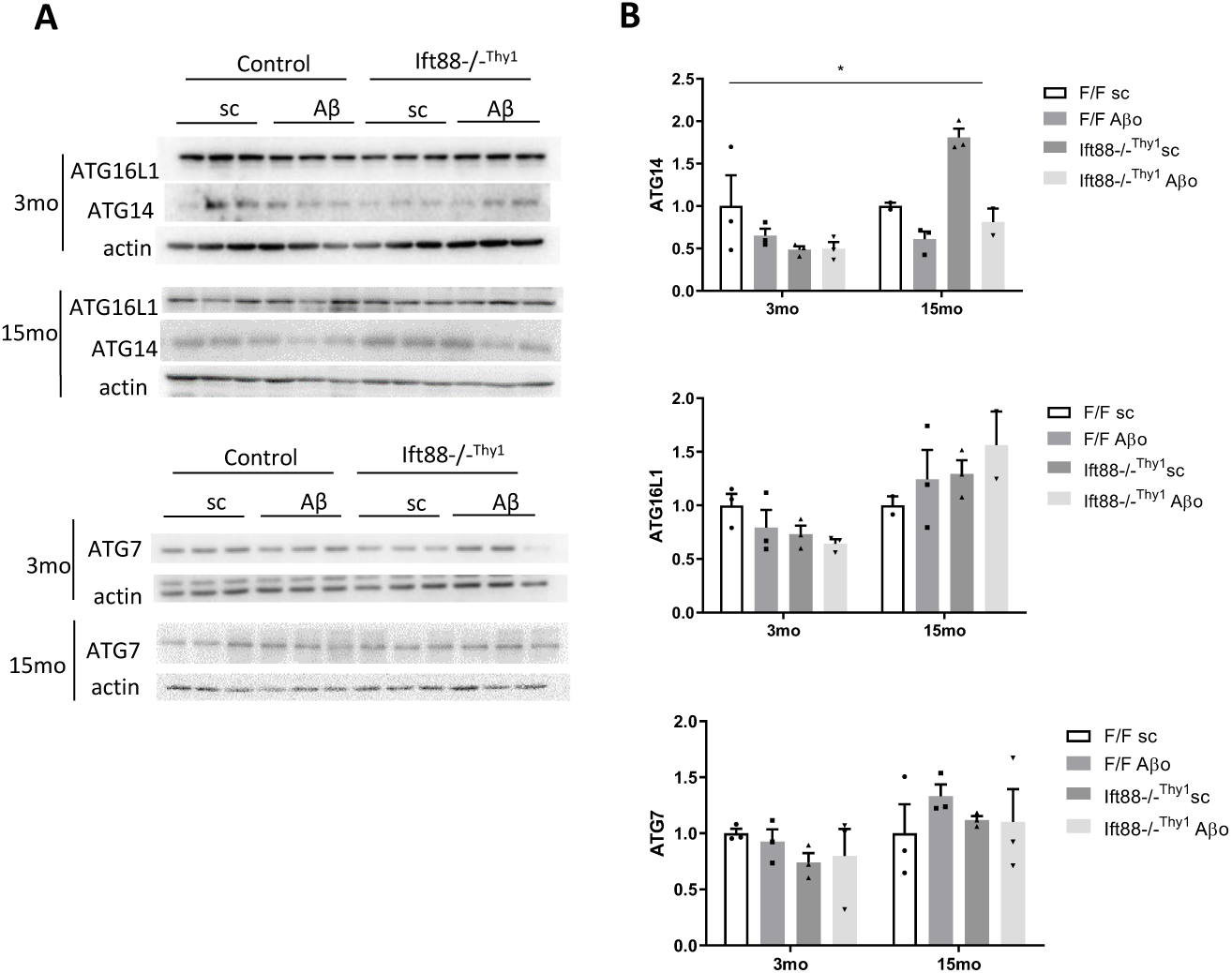
Early autophagy in female mice. (A) Immunoblots for early autophagy markers ATG16L1, ATG14, and ATG7, and actin in young (3mo) and old (15mo) F/F and IFT88-/-^Thy1^ female mice injected with either Aβsc or Aβo. (B) Densitometry of immunoblots in (A). *n* = 3. Mean ± s.e.m is shown. Two-way ANOVA and Sidak’s multiple comparisons test; * *p =* 0.05, ** *p =* 0.01, *** *p =* 0.001.

**Supplementary Figure 4.**
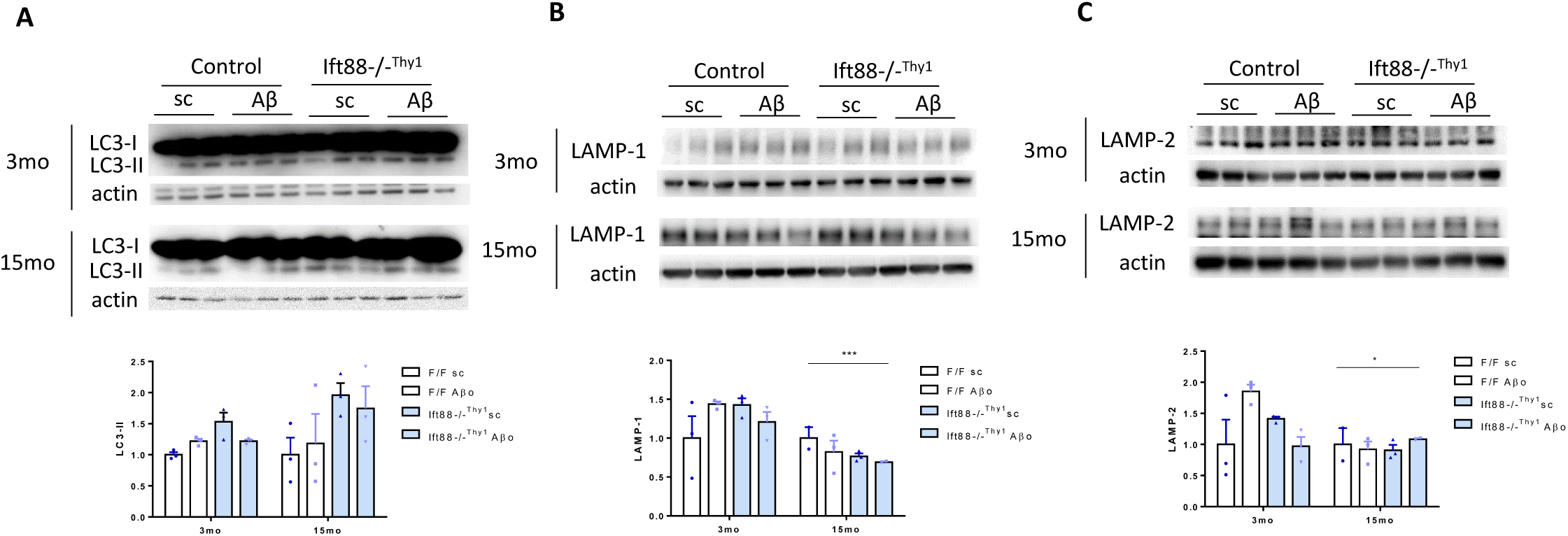
Autophagy in females. (A) Up; Immunoblot for LC3 and actin in young (3mo) and old (15mo) F/F and IFT88-/- ^Thy1^ female mice injected with either Aβsc or Aβo. Down; densitometry. (B) Up; Immunoblot for LAMP-1 and actin in young (3mo) and old (15mo) F/F and IFT88-/-^Thy1^ female mice injected with either Aβsc or Aβo. Down; densitometry. (C) Up; Immunoblot for LAMP-2 and actin in young (3mo) and old (15mo) F/F and IFT88-/-^Thy1^ female mice injected with either Aβsc or Aβo. Down; densitometry.

**Supplementary Figure 5.**
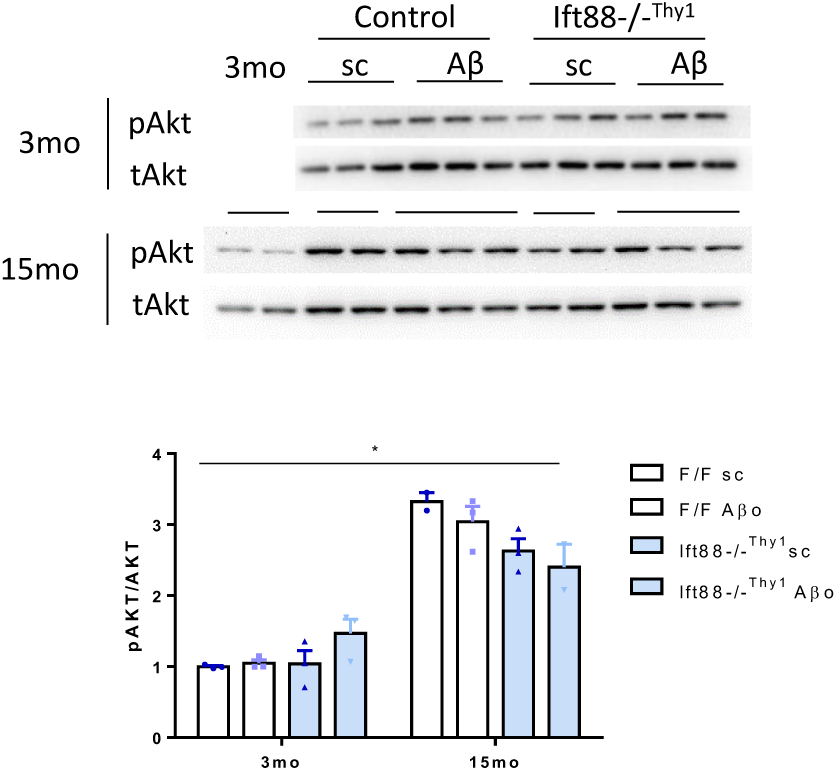
Autophagy regulation in females. Up; Immunoblot for phosphor-Akt and total Akt in hippocampus from F/F and IFT88-/-^Thy1^ fwmale mice injected with either Aβsc or Aβo. Down; Densitometry. *n* = 3. Mean ± s.e.m is shown. Two-way ANOVA and Sidak’s multiple comparisons test; * *p =* 0.05, ** *p =* 0.01, *** *p =* 0.001.

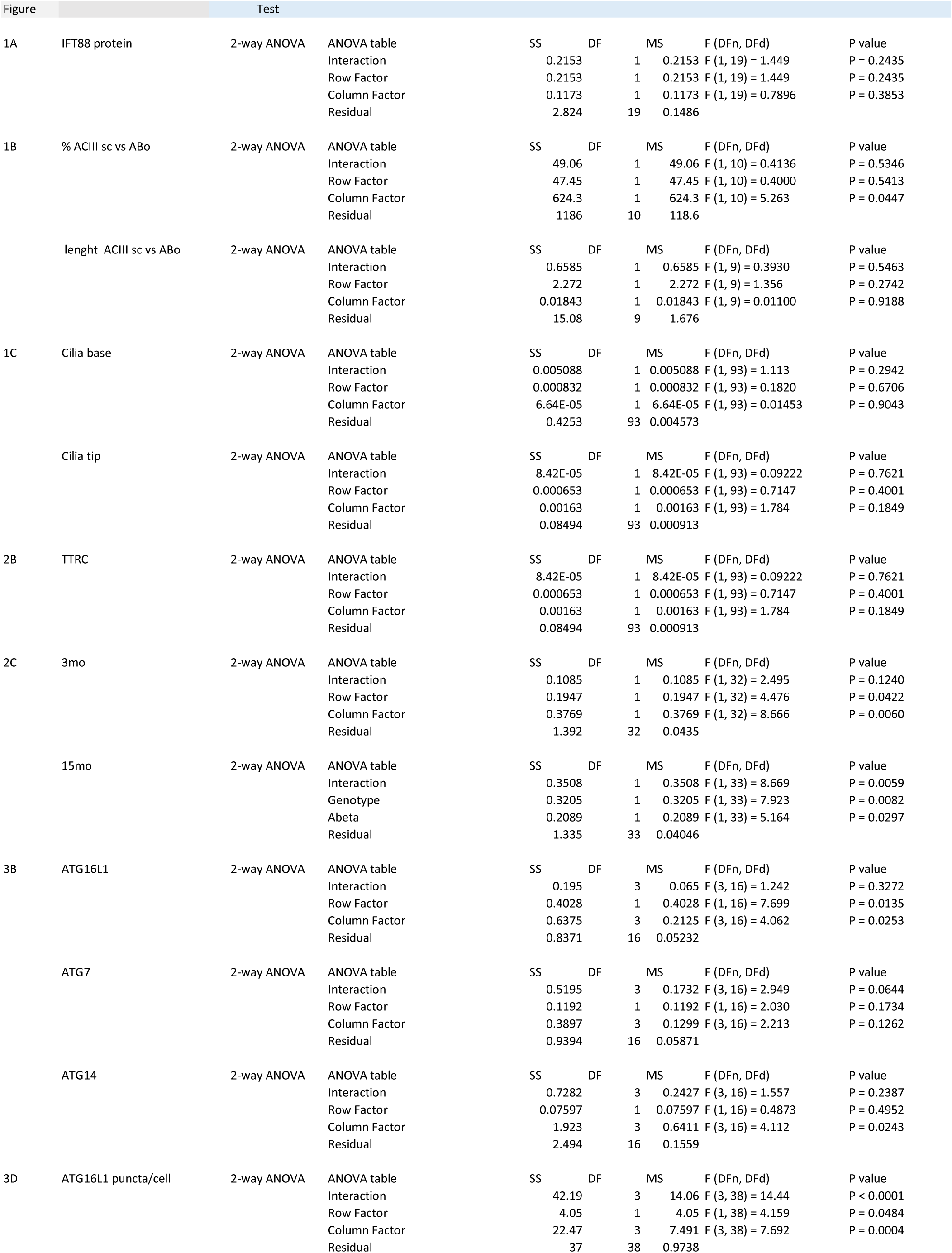
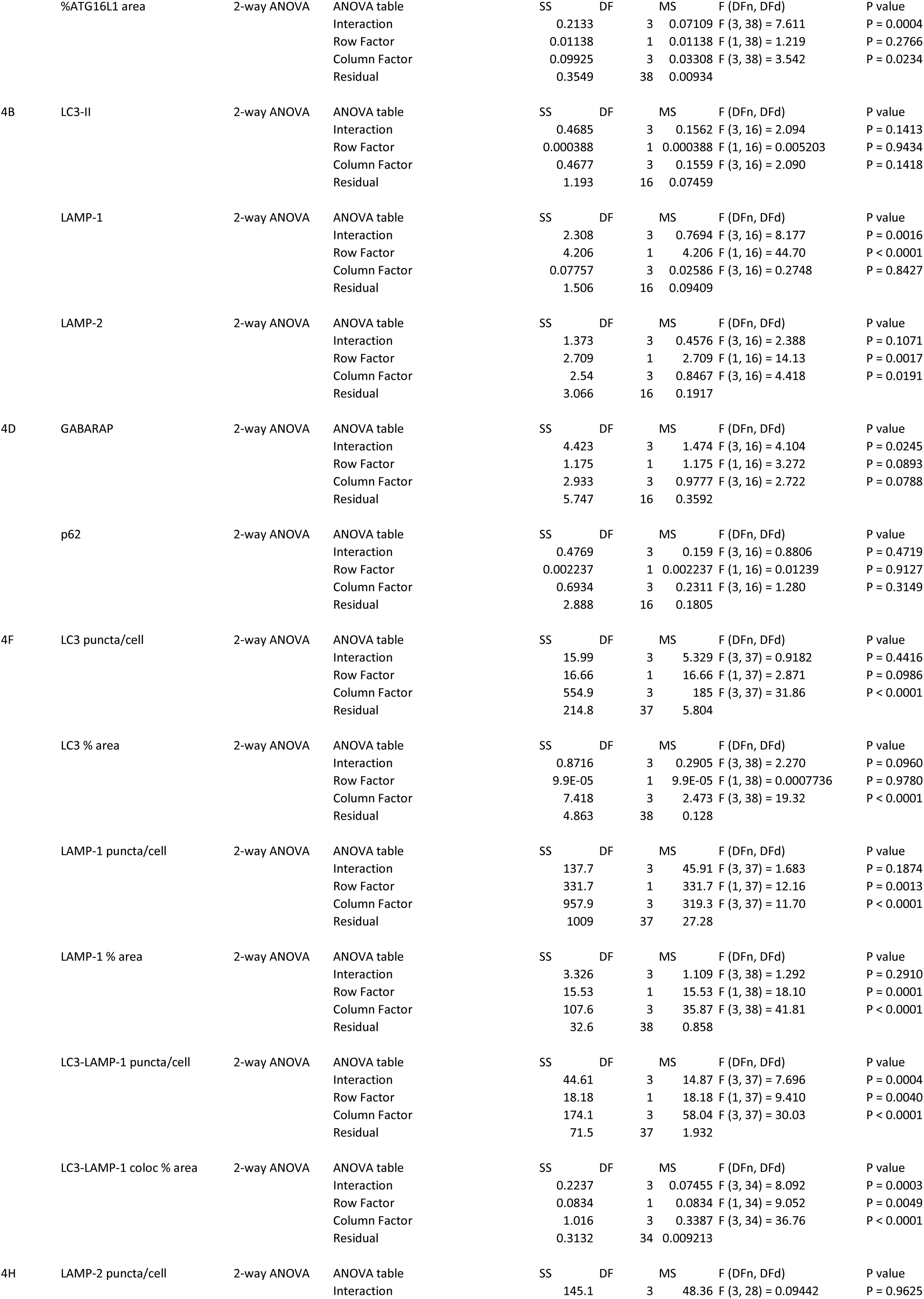
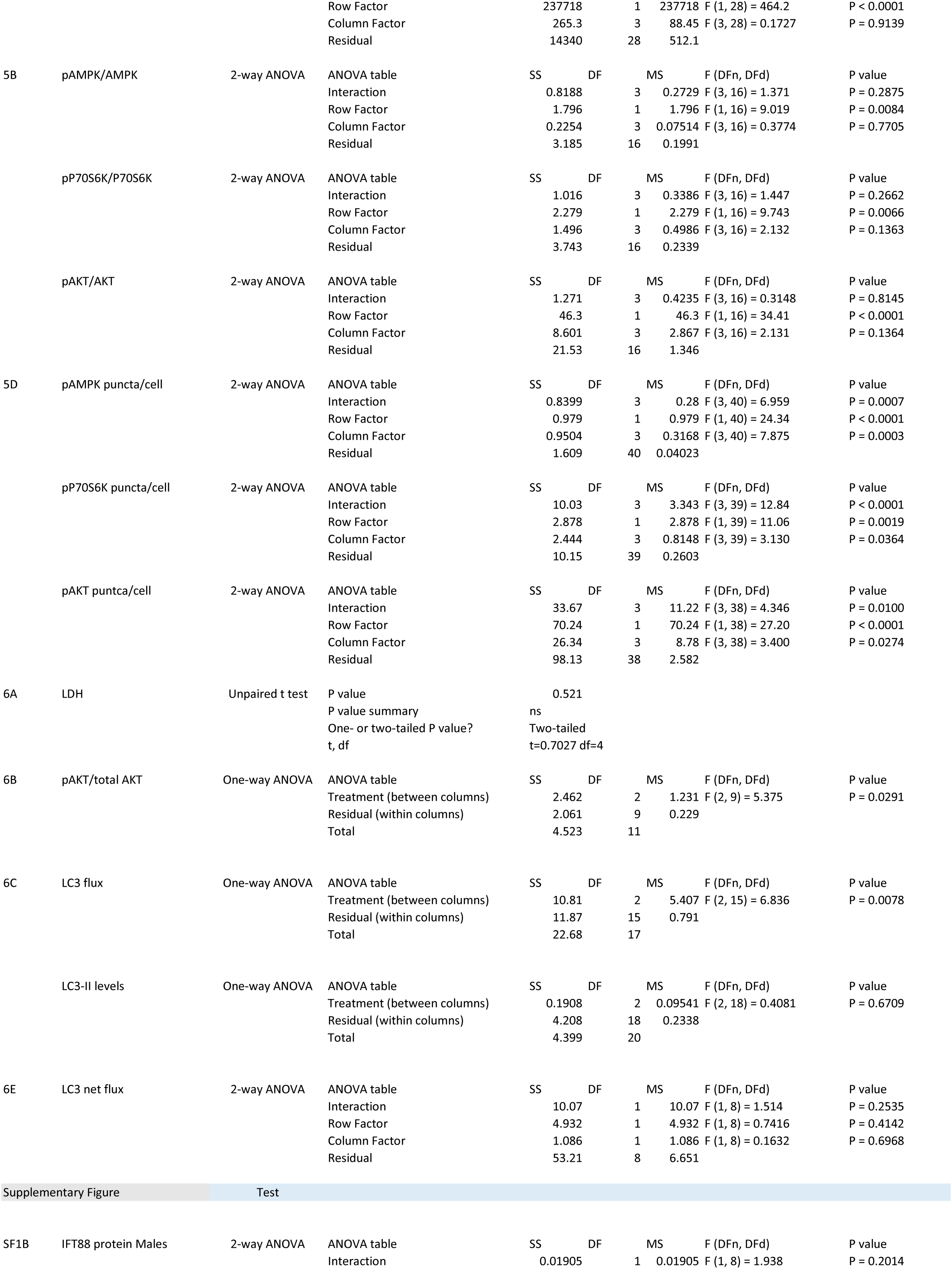
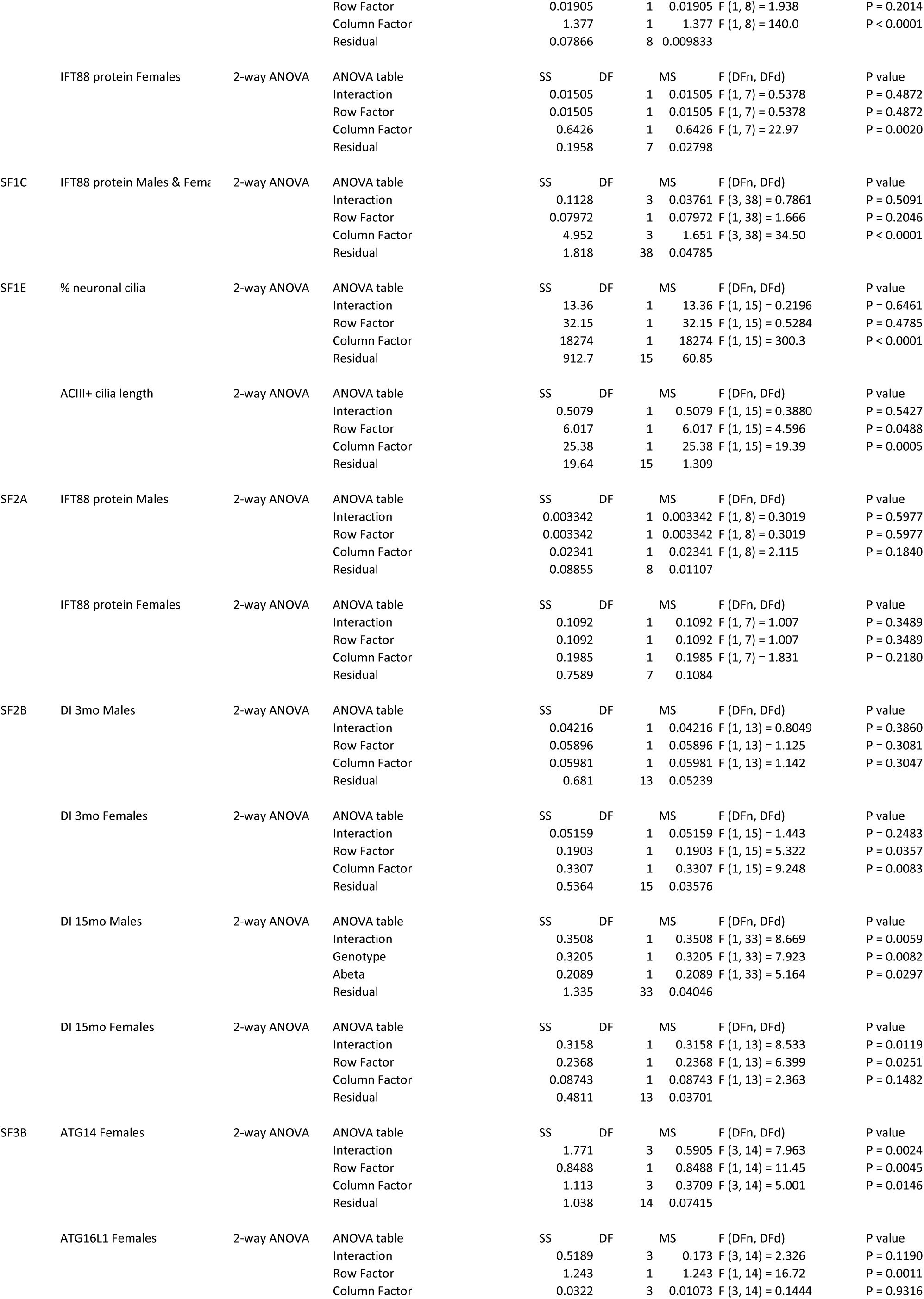
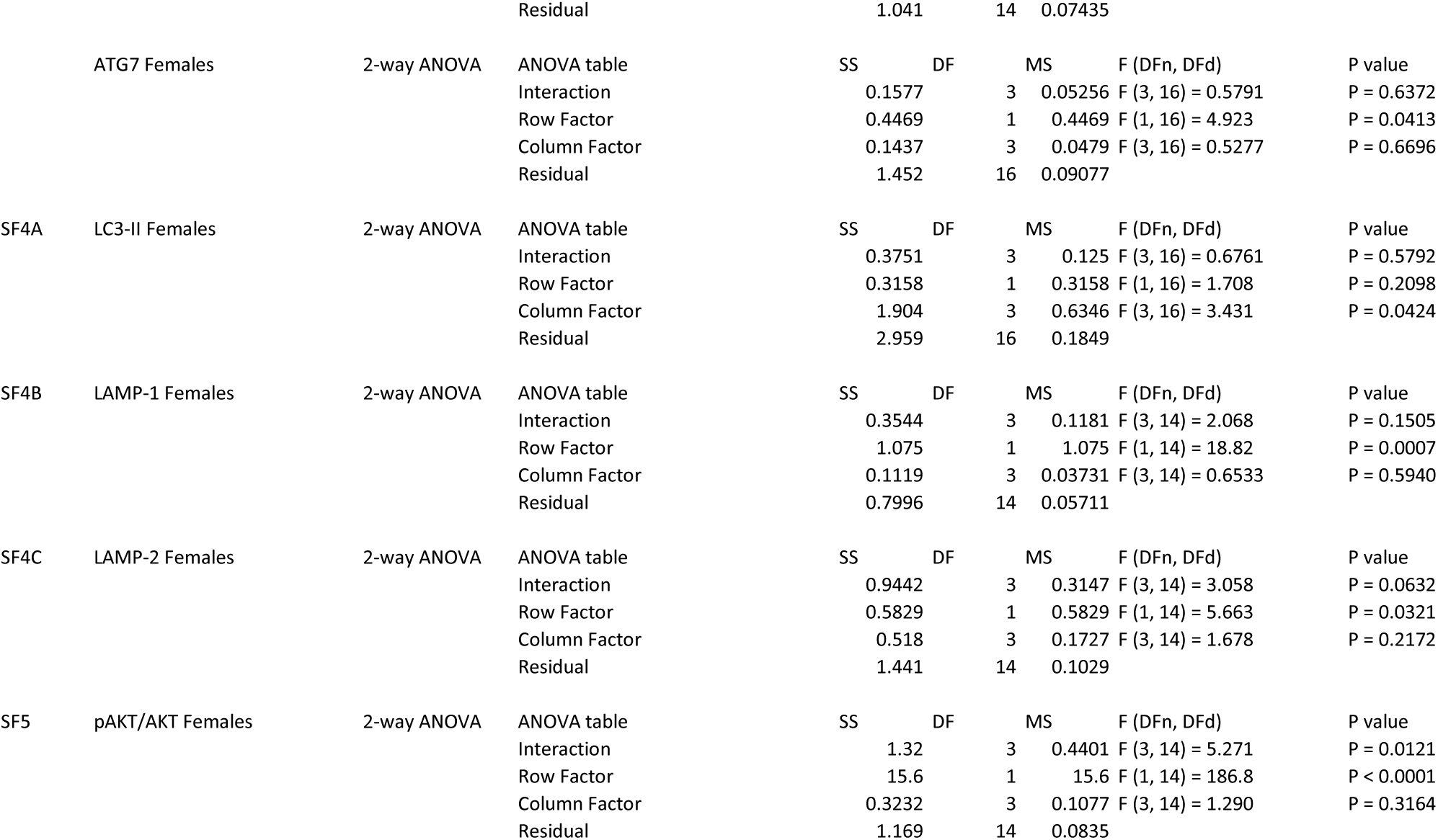

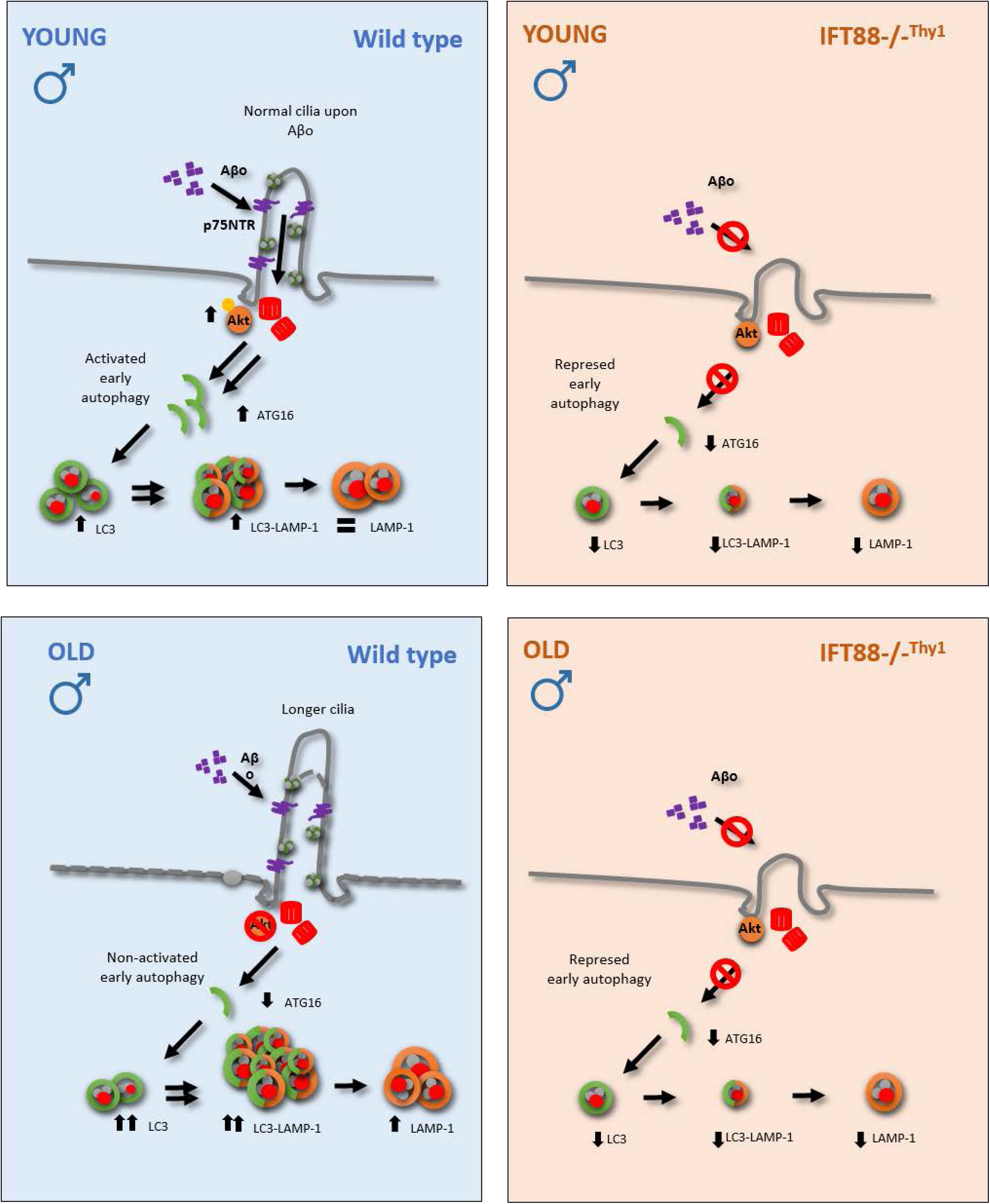

